# Detection of PatIent-Level distances from single cell genomics and pathomics data with Optimal Transport (PILOT)

**DOI:** 10.1101/2022.12.16.520739

**Authors:** Mehdi Joodaki, Mina Shaigan, Victor Parra, Roman D. Bülow, Christoph Kuppe, David L. Hölscher, Mingbo Cheng, James S. Nagai, Michaël Goedertier, Nassim Bouteldja, Vladimir Tesar, Jonathan Barratt, Ian S.D. Roberts, Rosanna Coppo, Rafael Kramann, Peter Boor, Ivan G. Costa

## Abstract

Although clinical applications represent the next challenge in single-cell genomics and digital pathology, we still lack computational methods to analyze single-cell and pathomics data to find sample level trajectories or clusters associated with diseases. This remains challenging as single-cell/pathomics data are multi-scale, i.e., a sample is represented by clusters of cells/structures and samples cannot be easily compared with each other. Here we propose PatIent Level analysis with Optimal Transport (PILOT). PILOT uses optimal transport to compute the Wasserstein distance between two individual single-cell samples. This allows us to perform unsupervised analysis at the sample level and uncover trajectories or cellular clusters associated with disease progression. We evaluate PILOT and competing approaches in single-cell genomics and pathomics studies involving various human diseases with up to 600 samples/patients and millions of cells or tissue structures. Our results demonstrate that PILOT detects disease-associated samples from large and complex single-cell and pathomics data. Moreover, PILOT provides a statistical approach to delineate non-linear changes in cell populations, gene expression, and tissue structures related to the disease trajectories supporting interpretation of predictions.

## Introduction

Single-cell genomics and digital pathology methods are revolutionary technologies, which in principle, allow researchers to computationally dissect molecular, cellular, and structural changes in human tissues^1,2^. However, the clinical application of single-cell sequencing, i.e., finding cells and their markers for patient stratification and personalized treatments^3^, is still in its infancy. Recent clinical genomics efforts include the use of single-cell transcriptomics to dissect the progression of acute human myocardial infarction^4^, to characterize COVID-19 patients’ disease severity^5^ and the development of pancreatic ductal adenocarcinomas^6^, just to cite a few. Computational analysis of these datasets mainly leverages standard methods for standard single-cell sequencing analysis, i.e., finding genes with differential expression in control vs. disease for each cell cluster. These approaches require a-priori patient classifications, i.e., control vs. disease. Therefore, they cannot be used to find novel sub-groups of patients. Alternatively, trajectory analysis can be performed to uncover disease progression allowing the characterization of early disease events. Particularly challenging is the multi-scale nature of single-cell experiments, i.e., each sample is represented by thousands of single cells, which are clustered in distinct cell types or cell states. Currently, there are no analytical methods to compare two single cell experiments from the same tissue from two distinct individuals.

Pathomics, i.e., the use of machine learning methods to extract morphological structures in histology slides, is likely to revolutionize pathology by producing reproducible and fast quantification of histological slides for disease diagnosis, as recently shown in the kidney^7^. Pathomics data of a slide is represented by thousands of anatomical structures, which are described by morphometric features, e.g., their individual shape and size. These can be clusters to find structures at distinct morphological states, i.e. distinct degrees of dysmorphism. As for scRNA-seq, there is no analytical method which is able to compare two or more histological slides based on morphometric properties of their structures.

Until now, only a few methods allow the analysis of single-cell genomics datasets at a sample level. PhEMD^8^ is based on earth moving distance (EMD) to measure the distance between specimens (single-cell samples), where the distance between specimens was based on clustering representations from a diffusion-based space. This method was successfully used to explore the response of cell lines to drug effects. Nevertheless, it is based on the diffusion map and pseudotime estimates on the cell level. This explicitly assumes the presence of a cellular continuum (cell differentiation/activation) between all cells in the scRNA-seq experiments. Therefore, it is not suitable for the analysis of single-cell experiments measured in whole organ samples with heterogeneous cell populations. Moreover, PhEMD lacks methods for interpretation of the results, i.e., detection of molecular and cellular features explaining predictions. Ravindra and colleagues ^9^ propose the use of graph attention networks for the classification of scRNA-seq samples and apply this to predict the disease states of multiple Sclerosis^10^. More recently, SCANCell^11^, which explores association networks across cell clusters, was proposed and applied for the analysis of systemic lupus erythematodes. Multiscale PHATE^12^ explores non-linear embeddings and multi-resolution representation to find the individual sample representation. This method uses supervised filters to find groups of cells related to COVID-19 mortality. Recently, Flores and colleagues proposed a multi-omics factor analysis that analyses single cell data at pseudo-bulk level^13^. This work focus on finding factors and molecular features explaining previously known sample phenotypes. Except for PhEMD, all related methods^9,11,12^ require labels of patients for their analysis and cannot be used in the unsupervised analysis (clustering or trajectory inference) of patient-level single-cell experiments.

## Results

### Patient level distance with Optimal Transport (PILOT)

Upon diseases, tissues undergo cellular and tissue remodelling changes. For example, in myocardial infarction cardiomyocytes acquire an injury cell states, immune cells migrate to injured tissue and fibrosis or scarring takes place to compensate tissue loss due to necrosis. We hypothesize therefore that changes in cellular composition are hallmarks of disease progression. We propose PILOT— Patient level distance with Optimal Transport (OT)— to find similarities between samples measured with multi-scale single-cell or pathomics data with the optimal transport based Wasserstein distance^14^ (Fig. 1). PILOT models each sample/patient as a distribution of cells into clusters (cell types and cell states present in a given tissue). It then uses optimal transport to move cell distributions between all samples. PILOT explores the fact that some cellular changes, i.e. the change of a healthy cardiomyocytes to injured cardiomyocyte state are short term events in disease progression, while tissue remodeling (e.g. replacement of lost cardiomyocytes by fibroblasts due to scarring process) is indicative of a long term event in disease progression. Therefore, we define a cost matrix for the optimal transport so that transporting masses between similar cell types (healthy vs. injured cardiomyocytes) have a lower cost than transporting masses between distinct cell types (cardiomyocytes vs. fibroblasts). The optimal transport plan can be used to obtain the Wasserstein distance. PILOT explores diffusion maps^15^ and a path inference algorithm^16^ to infer sample-level disease progression trajectories or graph-based clustering^17^ to find unknown sub-groups of patients. For disease trajectories, PILOT explores robust non-linear and regression models^18^ to find cell types, genes, or morphological features associated with disease progression. The test indicates for features explaining the predicted pseudotime by fitting either linear or quadratic models, i.e. find changes in cell populations. For the case of genes and morphological features, the test can be conditioned on a cluster/cell type. Moreover, we use a Wald test to check if the cluster specific model fit differs from a global model^19^. Thus PILOT represents the first approach to detect unknown patient trajectories and clusters, while providing statistical models for interpretation of cellular, molecular, and morphometric features associated with disease progression.

**Figure 1.**
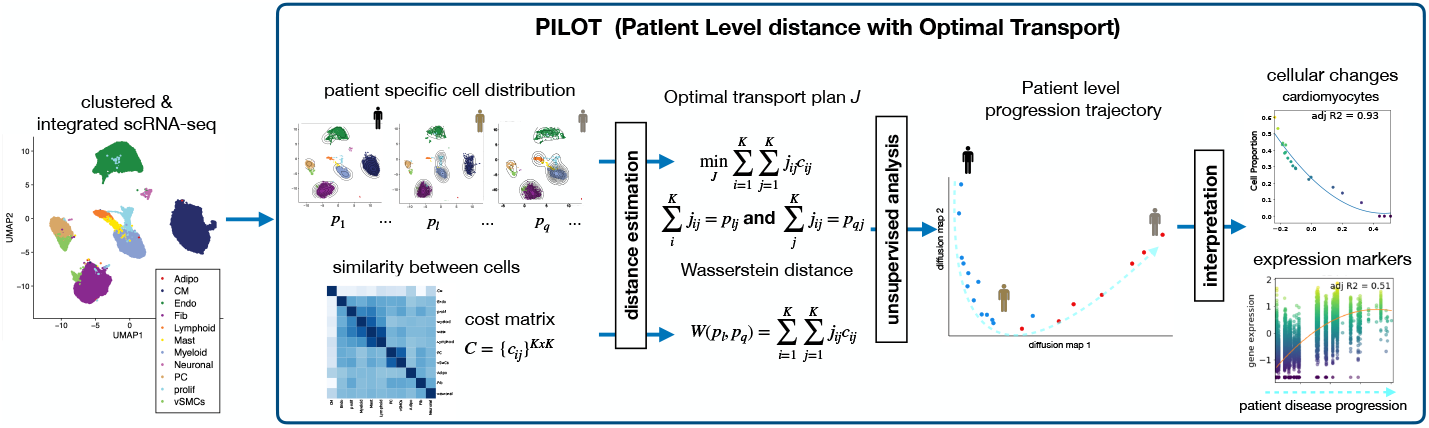
PILOT schematic and benchmarking: **A**. PILOT receives as input clustered and integrated scRNA-seq data. Next, it estimates patient-specific cluster distribution and a cost function (similarity between clusters). These are used as input for an optimal transport algorithm^14^. The transport costs provide a distance (Wasserstein), which can be used for unsupervised analysis, such as a diffusion map and disease progression analysis. Finally, PILOT uses non-linear regression to find cellular, molecular, or morphological features associated with disease progression.

### Evaluation of patient-level clustering and trajectory analysis

We compare the results of PILOT and PhEMD^8^ and baseline methods in the recovery of known patient groups of eight single-cell RNA data sets (PBMCs from systemic lupus erythematodes^20^, pancreatic ductal adenocarcinoma [PADC]^6^, acute myocardial infarction^4^, PBMC of patients with COVID-19 infection^5^, kidney injury^21^, lung cancer^22^, follicular lymphoma^23^ and diabetes^24^) and four pathomics data sets with morphological features of glomeruli and tubules from two distinct disease cohorts (kidney IgA nephropathy of the VALIGA study [IgAN]^25^ and kidney biopsies of the Aachen cohort [AC]^7^). For most of these data, we use disease status (control and case) as labels for the clustering task. For the Kidney IgAN pathomics data, patients/donors were labeled by their estimated glomerular filtration rate (eGFR; normal, reduced, and low); for the COVID-19 samples were labeled as control, mild and severe cases; for the scRNA-seq kidney injury data samples were grouped as normal, chronic kidney disease and acute kidney failure. The lung and pancreas cell atlas have controls and distinct disease types (4 for lung and 3 for pancreas). These are the largest publicly available single cell and pathomics data with up to millions of cells and structures and measured in hundreds of patient samples (Table 1). We also evaluate two baseline methods: pseudo-bulk libraries per sample by using state-of-art RNA-seq pipelines^26^ or clustering directly the cell proportion matrices.

**Table 1.** Characteristics of data sets used for benchmarking.

| Data | Type | #Cells | #Samples | #Cell_types | #Classes |
| --- | --- | --- | --- | --- | --- |
| COVID-19 PBMC | scRNA-seq | 993,171 | 151 | 10 | 3 |
| Myocardial Infraction | scRNA-seq | 115,517 | 20 | 11 | 2 |
| Lupus PBMC | scRNA-seq | 1,263,676 | 261 | 11 | 2 |
| Pancreas (PDAC) | scRNA-seq | 57,530 | 35 | 10 | 2 |
| Kidney | scRNA-seq | 76,020 | 36 | 57 | 3 |
| Lung | scRNA-seq | 941,504 | 165 | 33 | 5 |
| Diabetes | scRNA-seq | 264,235 | 52 | 13 | 4 |
| Follicular lymphoma | scRNA-seq | 137,147 | 23 | 18 | 2 |
| Kidney IgAN (Glomeruli) | Pathomics | 24,227 | 634 | 14 | 3 |
| Kidney IgAN (Tubule) | Pathomics | 64,493 | 634 | 15 | 3 |
| Kidney AC (Glomeruli) | Pathomics | 4,731 | 57 | 28 | 2 |
| Kidney AC (Tubule) | Pathomics | 56,998 | 57 | 7 | 2 |

For scRNA-seq data, we used the data as provided in the original manuscript as input for PILOT. For pathomics data, structure segmentation was performed with FLASH followed by graph-based clustering as described before^7^. Next, we used graph-based clustering^17^ on the Wasserstein distance matrices and evaluated results with the adjusted Rand index (ARI)^27^. The number of clusters was the same as the number of true classes in the data. PILOT obtained the highest average ARI score and its ranking was significantly superior than all competing methods (Fig. 2A, *p*-value < 0.05; Friedman-Nemenyi test, and Sup. Table S1). Next, we evaluate the performance of distance matrix of PILOT and competing methods with the Silhouette index^28^. PILOT had the highest ranking among all evaluated methods (Fig. 2B) and outperformed PheEMED (*p*-value < 0.05; Friedman-Nemenyi test). Due to the large size of data sets, computing time is also a relevant aspect. In the Lupus PBMC data with millions of cells, PILOT required 58.12 seconds versus 9.03 seconds of the simple pseudo-bulk and 16.50 minutes of PhEMD ^1^. These indicate that PILOT and some competing methods can be efficiently run in large data with millions of cells or tissue structures.

**Figure 2.**
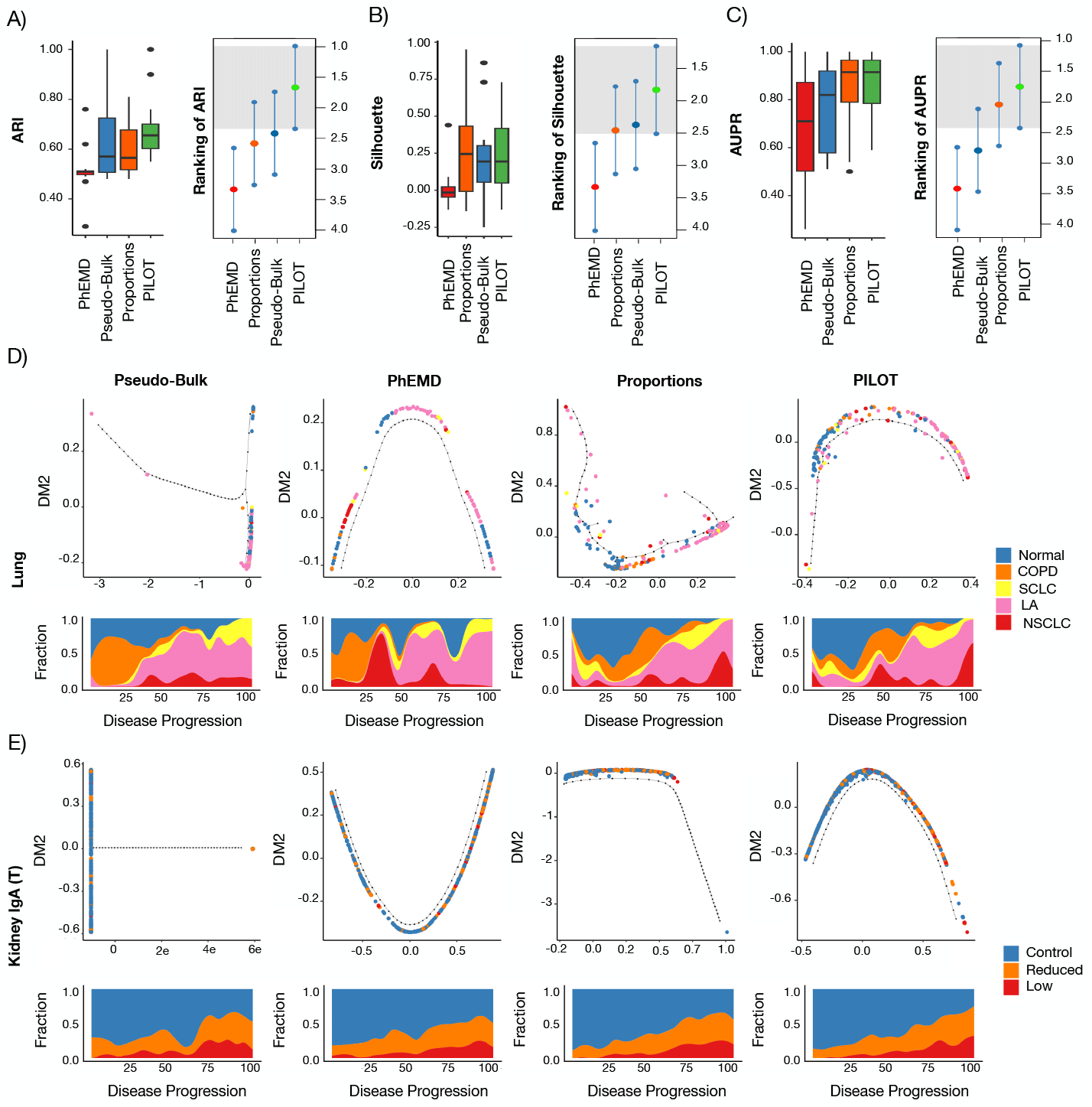
**A)** Box plot with ARI values (y-axis) for distinct evaluated methods (x-axis) (left) and ranking distributions (x-axis) based on the Friedman-Nemenyi test for distinct methods (right). Methods with average ranking in the gray area are top performers. **B)** and **C)** are equivalent of **A** for Silhouette and AUPR statistics. **D)** Difusion maps (top) and fraction of patient labels vs. pseudotime (bottom) on lung cell atlas data set for distinct algorithms. For this data, labels COPD, SCLC, LA, and NSCLC corresponds to Chronic Obstructive Pulmonary Disease, Squamous Cell Lung Carcinoma, Lung Adenocarcinoma, and Non Small Cell Lung Carcinoma groups respectively. **E)** Difusion maps (top) and fraction of patient labels vs. pseudotime (bottom) for Kidney IgAN (Tubule).

We observe that diffusion maps estimated with PILOT recovered trajectory-like structures in all analyzed data sets (Fig.2D-E; Sup. Fig. S1-S2). Therefore, we modeled these trajectories with the EIPGraph^16^ to rank samples regarding their disease progression score. We observe that a disease progression score recapitulates the severity of diseases in the data sets (Fig. 2D-E; Sup. Fig. S3).. For example, in the lung cell atlas normal samples were at the beginning of the trajectory. These were followed by samples associated with chronic lung disease (COPD), while acute carcinoma samples (SCLC, LA, NSCLC) accumutated in the end of the trajectory. For Kidney IgAN (Tubule), we observe the trajectory order samples regarding normal, low or reduced glomerular filtration rate. To evaluate this more systematically, we considered the two-class problem for all data sets (controls vs. disease) and used the estimated pseudo-time score to compute the area under the precision-recall curves (AUPR). We observe that PILOT has the highest mean rank when considering AUPR values, which supports its power in disease trajectory detection (Fig. 2, *p*-value < 0.01; Friedman-Nemenyi test).

PILOT requires no definition of parameters. However, it assumes that single cell experiments have been previously preprocessed and clustered. Currently, PILOT adopts the same clustering and integration strategy as in the manuscript describing the data, as cells are labelled and this helps the interpretation of results. To investigate if clustering can impact PILOT, we change the resolution parameter of the Leiden Clustering algorithm for selected data sets. We observe that this parameter is not critical for the performance of PILOT (Sup. Fig. S4).

### PILOT trajectories detects events associated to cardiac remodelling in myocardial infarction

Next, we analyze the inferred trajectory from normal heart tissue towards ischaemic zone (IZ) tissue in acute myocardial infarction. The trajectory reflected the known annotation of samples with the exception of one case in the decision boundary between controls/IZ samples (Fig. 3A). We use non-linear regression methods to find cellular and molecular changes associated with the disease progression score. Major cellular changes, as indicated by the highest adjusted *R*^2^ values, include a quadratic increase of SPP1 positive macrophages and myofibroblasts, a quadratic decrease of healthy cardiomyocytes followed by a later and smoother decrease of stressed cardiomyocytes during disease progression (Fig. 3B; Sup. Fig. S5). These patterns are in accordance with cellular changes in early myocardial infarction, which include damage of myogenic tissue followed by inflammation and fibrosis. This example also confirms the power of PILOT predicting trajectories to find continuous changes related to tissue remodelling.

**Figure 3.**
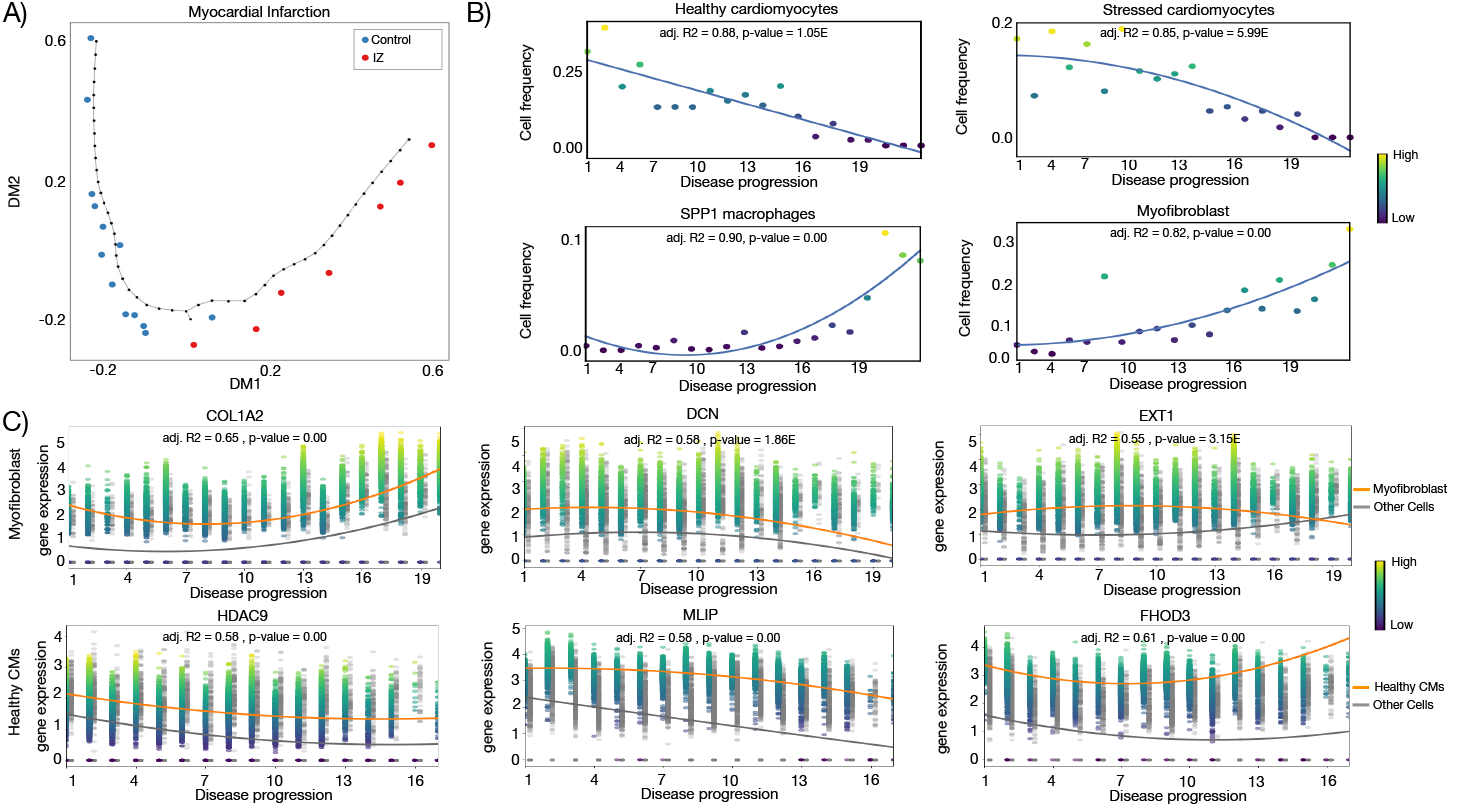
Sample level analysis of myocardial infarction: **A**. Diffusion map and disease progression trajectory in myocardial infarction. **B**. Fraction (y-axis) of top four cells (highest R2) vs. disease progression (x-axis). **C**. Expression (y-axis) of selected genes with significant changes in disease progression (x-axis) for healthy cardiomyocytes and myofibroblast cells.

At the gene level, PILOT identifies genes related to extracellular matrix remodelling induced by myofibroblasts (Supp. Fig. S6). These include COL1A2, which has an overall quadratic increase in gene expression upon disease progression (Fig. 3C). Decorin (DCN) and exostosin-1 (EXT1) are examples of genes with decreased expression upon late stages of disease progression. Decorin is a fibroblast specific gene with antifibrotic properties due to TGFB signalling inhibition^29,30^. Exostosin-1 has been associated with early formation of collagen fibres^31^. We observe genes associated with heart development and with disease progression related changes in healthy cardiomyocytes (Supp. Fig. S8). One of these genes with a steep reduction in expression is HDAC9, which functions as a signal-responsive repressor of cardiac hypertrophy by inactivating MEF2 transcription factors^32^. MLP, which is a gene with a slow decrease in expression, is a regulator of myofibril assembly ^33^. Finally, FHOD3 is a gene important in the regulation actin filaments. It has a decrease in expression at sames at the middle of the trajectory indicating a potential role which would have been missed if not analyzed within temporal context^34^. Altogether, these results support how PILOT can detect molecular and cellular changes at distinct stages of myocardial infarction.

### PILOT trajectories in Pathomics data

Next, we performed a trajectory analysis of the pathomics data of kidney IgAN biopsies. As distinct morphological features are used to describe tubules and glomeruli, it is not possible to cluster these structures together, so these data are independently analyzed as in^7^. We combine the two disease progression scores with their sum, which yields similar or higher AUCPR scores than the trajectories considering one structure at a time (Fig. 4A; Fig. S9). This trajectory is associated with a linear decrease in eGFR (Fig. 4B), which is the current surrogate used in clinical practice to estimate kidney function. Regarding morphometric features, PILOT detects a significant linear decrease of glomerular tuft sizes and tubule sizes; and a linear increase in distance between glomeruli and tubules as significant characteristics of disease progression (Fig. 4C; Supp. Fig. S10-S11). These are indicative of interstitial fibrosis, tubular atrophy, and glomerulosclerosis, which are all hallmarks of kidney function decline. Finally, we made use of a prognostic variable from the kidney IgAN biopsies, which indicates if patients progressed to kidney failure. PILOT progression score is more associated with kidney failure (*p*-value of 2.4e-11; likelihood ratio test) than a multivariate model combining all morphometric variables (*p*-value of 7e-05; likelihood ratio test) or the use of individual morphometric variables (Supp. Table S2). Indeed, samples with higher disease progression scores (top 75 quartiles) have a higher chance of kidney failure than other samples (Fig. 4D). Altogether, these results reinforce the power of PILOT in discriminating disease outcomes from pathomics data.

**Figure 4.**
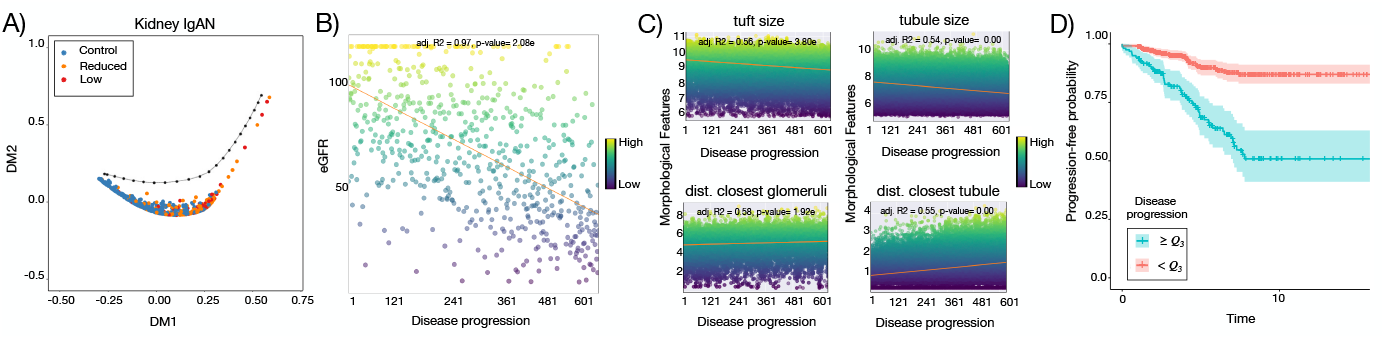
Sample level analysis of kidney IgAN : **A**. Diffusion map and disease progression trajectory in kidney IgAN. **B**. GFR values (y-axis) verhttps://www.overleaf.com/project/643d10d6312455328d1029d9sus disease progression (x-axis). **C**. Morphometric values (y-axis) of kidney structures vs. disease progression (x-axis). **D**. Kaplan-meyer plot with the time to kidney failure from patients within the upper quartile (high progression score) vs. other samples (low progression score).

## Discussion

The technological improvements in single cell genomics and digital pathology are providing us with clinically rich and large-sized data describing cellular and morphological changes in diseases. We present here PILOT—the first computational pipeline for the detection and feature characterization of disease trajectories from single-cell genomics or pathomics data. By using a comprehensive benchmark with twelve data sets, we show that PILOT is superior to the competing approaches in both clustering and disease trajectory prediction. Another important aspect, which is only addressed by PILOT, is the interpretation of predictions. PILOT uses a robust non-linear regression model, which can predict cellular populations, marker genes or morphological features associated with the disease progression.

We revisited the analysis of the largest single-cell genomics data on myocardial infarction^4^, where PILOT could successfully predict a trajectory from controls towards ischemic samples. This allowed us to find non-linear changes in cell composition during cardiac remodeling, i.e., quadratic decrease of healthy cardiomyocytes and quadratic increase of myofibroblast and macrophages cells. Similarly, PILOT dissected gene expression programs associated with these changes, such as non-linear increase in expression of extra cellular matrix related genes in myofibroblasts and increase of genes associated with cardiomyopathy in healthy cardiomyocytes. We also evaluated the power of PILOT in inferring a disease trajectory of pathomics data from patients with kidney IgAN. We show that disease progression as estimated by PILOT is related to eGFR and provides a better predictor for future kidney failure than the use of the morphometric features alone or together. These highlight the power of PILOT in finding non-linear manifolds to model disease progression. Molecular studies of diseases are increasingly based on multi-modal measurements; gene expression, protein abundances, chromatin accessibility, and histology images at either single cell and/or spatial level^35^. Future work of multi-scale level analysis will require methods considering the multi-modal and spatial nature of these data into account.

## Methods

### PILOT

We formalize the main steps of PILOT^2^. In analyzing single-cell experiments, assume there are *L* single-cell matrices 𝒳 = {**X**_1_, …, **X**_*l*_, …, **X**_*L*_}, where 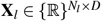. Where *N*_*l*_ represents the number of cells in sample *l* and *D* is the number of features (genes), which is common between all experiments. In practice, PILOT receives as input a single matrix **X**^**I**^ ∈ {ℝ}^*N*×*D*^ after integration of the matrices in 𝒳, where *N* represents the total number of cells. To keep the sample information, we define a vector indicating the sample identity of cell *i*, i.e. *s* = {*s*_1_, …, *s*_*N*_}, where *s*_*i*_ ∈ {1,…, *L*} indicates which sample (patient) the cells belongs to. A usual representation/sumarization of single cell experiments is to group cells with a clustering algorithm. This can be represented by a vector *y* = *y*_1_, …, *y*_*N*_, where *y*_*i*_ ∈ {1,…, *K*} indicates the group of cell *i*.

Our main problem is to estimate the distance between scRNA-seq data measured over two distinct samples (patients), where a scRNA-seq is represented as a set of clustered cells. Here we explore the concepts of the optimal transport based Wasserstein method^36^ to compare two samples by representing samples as distributions of cells. First, we defined the probability of having a cluster *k* for patient *l* as:

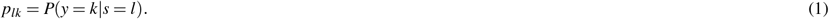

This can be used to define a probability distribution vector *p*_*l*_ = (*p*_*l*1_, …, *p*_*lK*_).

From these, we use optimal transport to find the optimal transport plan *J* = {*j*_*ij*_}^*KxK*^ between two distributions *p*_*l*_ and *p*_*q*_ describing samples *l* and *q* by considering that there is a cost *c*_*ij*_ associated with moving some mass between cluster *i* to cluster *j*:

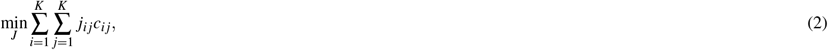

such that *j*_*ij*_ ≥ 0, 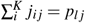 and 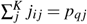 and 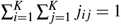.

For a given optimal transport plan *J*, the Wasserstein distance (*W*) is calculated as:

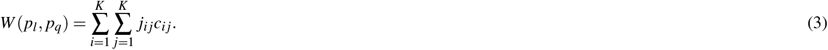

The optimal transport matrix *J* can be estimated using a minimum flow cost algorithm^36^. PILOT is based on the emd2 function from POT library^37^, which implements the best solver described in^36^. By estimating the Wasserstein distance between all pairs of samples, we obtain a distance matrix *W* between all samples.

This framework, which is also denoted Earth Moving Distance^38^, is also explored by PhEMD. PILOT, however, explores important issues which are crucial in the noisy and large nature of single cell and pathomics data. These are namely: (1) how to obtain robust estimates of probability distributions due to potential cell content bias and low cell coverage of individual samples; (2) how to estimate the cost matrix *C* on the large matrices **X**^**T**^; and (3) by offering statistical models to select features associated with disease progression. These three points are described below.

### Robust estimation of sample probability distributions

The coverage of cells per cluster can vary across distinct samples. This effect is potentially higher in diseased samples, due to their lower cell viability. Let *z*_*l*_ = (*z*_*l*1_, …*z*_*li*_, …, *z*_*lK*_) be a vector, where *z*_*lk*_ is the number of cells in a cluster *k* and sample *l*. We define the following hierarchical model:

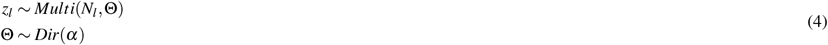

where *N*_*l*_ is the number of cells in the sample *l*, Θ is a random variable representing distributions *p* and *α* is the hyper-parameter of a Dirichlet distribution. The posterior distribution can be re-written as

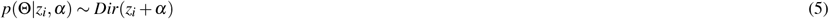

We can use this to obtain maximum a posteriori estimates of Θ, i.e.

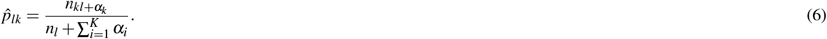

Here, we use the following parametrization of the prior *α*_*k*_ = *N*_*k*_*/N* ∗ *c*, where *c* is set as 0.1 as default. This prior adds “pseudo cell counts”, which are weighted by the cell distribution of all samples. This prior mitigates the fact rare cells might not be observed in samples with low cell coverage.

### Estimation of Cost Matrix

The optimal transport also considers the cost *c*_*ij*_ of transporting a distribution mass from a cluster *i* to a cluster *j*. For this, we generate a median representation for each cluster (in the PCA space) and estimate the cosine distance between the median value of clusters *i* and *j* as the cost to transport masses. The cosine distance between the median of cluster *i* (*M*_*i*_) and cluster *j* (*M*_*j*_) is defined as:

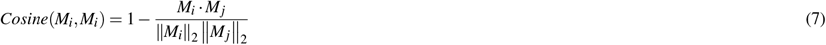

Of note, PhEMD uses centroids and Euclidean distances in non-linear embedding to estimate the cost matrices. The non-linear embedding assumes a cellular continuum between all cells in the data, which is not present in whole tissue single cell or pathomics data. Moreover, median values reduce the effects of outliers.

### Clustering and disease trajectory estimation

The matrix *W* (Eq. 3) provides the distance between all samples. Clustering analysis can be performed by providing *W* as input to a Leiden clustering algorithm^17^. PILOT also performs trajectory analysis by the use of diffusion maps^39^ followed by a trajectory estimation with EIPLGraph^16^. For this, we apply the Gaussian kernel to *W* ^40^ to construct the affinity matrix as *W*^*s*^, i.e.:

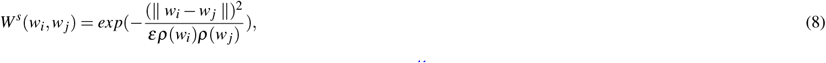

where *ε* and *ρ* are the scale parameter and the bandwidth function^41^ respectively.

Next, we compute the transition matrix:

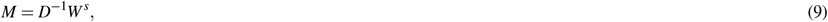

where *D* is a diagonal matrix with 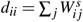. Finally, we perform the spectral decomposition of matrix *M* as:

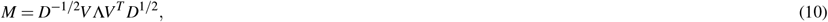

where Λ and *V* are the eigenvalue and eigenvectors matrices. Ultimately we use eigenvectors with the highest values for obtaining diffusion maps. These are provided as input for EIPLGraph^16^, which infers a backbone of the trajectory. It also allows us to rank samples with a disease progression score *t* = *t*_1_, …, *t*_*L*_, where *t*_*l*_ is the ranking of the sample *l*. In our experiments, we only considered two highest eigenvectors. EIPLGraph also requires a root sample, which was visually selected to reflect parts of the trajectory with control samples.

### Identification of molecular and structural features associated with disease progression

PILOT uses step-wise regression models to identify features (cellular abundances, gene expression or structural properties), whose values are consistently changing across disease trajectories. For this, it fits regression models with linear, quadratic, and linear-quadratic terms for each feature and uses statistical tests to determine the goodness of fit. We then report the most significant model for each feature.

Let *x*_*ij*_ represent the expression of a feature (gene) *j* in a cell *i*, and *p*_*i*_ the pseudo-time variable (associated with the pseudo-time of its sample, 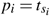). In short, we fit the three regression models to the *j*th feature:

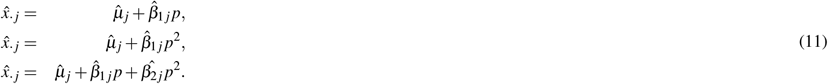

where 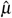 is the models’ intercept, and 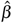 are the models’ coefficients. The models are ranked based on the coefficient of determination (*R*^2^), and we only consider the model with the highest R-squared and at a significance level (*p*-values < 0.05). The same approach can be performed for distinct features, such as the proportion of cell types in the sample or morphological features.

In the case of gene expression, one is mostly interested in finding cell type (cluster) specific genes. Therefore, we only consider cells *i* belonging to cluster *k*. Due to the sparsity of single cell data, i.e., dropout events, we observed a single side tailed distributions of residuals. Therefore, to improve the robustness of our regression models, we use the Huber least square criterion (Eq. 12)^42^.

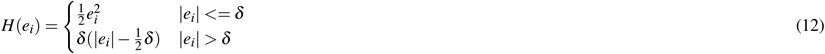

where 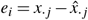. The Huber regression lessens the effects of the outliers by using a term *δ* (default of 1.35). This defines residual values associated with outliers. For large *δ*, the regression is equal to ridge regression.

Of note, the previous formulation requires a re-definition of the R-squared to consider the Huber regression penalization, that is:

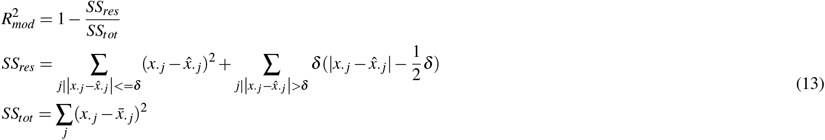

Another relevant questions is if the pattern of the feature (gene expression or morphometrics) in a given cluster over pseudo-time is distinct from other clusters. To do this, we compare the expression patterns along the pseudotime between one cluster vs cells/structures from all other clusters (non-cluster). We utilized Wald statistics, using the linear combination of coefficients, to test whether there are differences between pair points of fitted curves (cluster vs. non-cluster). Subsequently, for each gene, we test 2 × 3 null hypotheses (between two models, each having at most three coefficients; see Eq. 11) on the *n* pseudotime points:

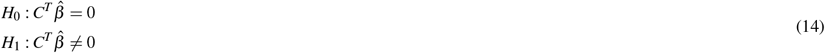

where *C* is the (2 × 3) × *n* contrast matrix of interest and 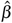 are coefficients of two models. The distribution of the test statistic under the null hypothesis is:

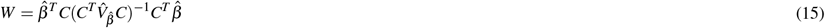

where 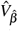 is an estimator of the variance-covariance matrix of estimated coefficients. For large enough *n, W* is distributed as *χ*^2^ with *r* degrees of freedom d.f. for *n* = 100^43^. To estimate the Wald score, we need to perform a eigendecomposition of the variance-covariance matrix 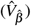. We consider all eigenvectors with eigenvalues larger than 1*e*^*-*8^. Statistical significance is estimated with the *χ*^2^ and the degree of freedom in the rank.

## Materials

### Data sets

We use public single cell and pathomics data sets to benchmark the proposed methods (see Fig.1B).

### Single cell data sets

Peng and colleagues^6^ performed a single cell study on pancreatic ductal adenocarcinoma (PDAC). They characterized 11 healthy and 24 PDAC samples with a total of 57,530 scRNA-seq annotated in 10 major pancreas cell types. As only raw data was provided (obtained from GSA: CRA001160 https://ngdc.cncb.ac.cn/gsa/browse/CRA001160), we reanalyzed the data using Seurat and re-annotated cell types using the same marker genes (Sup. Fig. S12).

Systemic lupus erythematosus is a common type of lupus that, in the immune system, pounds its tissues, yielding overall rash and tissue injury in the acted organs^20^. This single cell study characterized peripheral blood mononuclear cells (PBMCs) gene expression of 261 donors consisting of 1,263,676 cells. Of these donors, 99 were healthy controls, and 162 were disease patients. We use the normalized and clustered (11 groups) data sets provided in Gene Expression Omibus (GEO; GSE174188; GSE174188_CLUES1_adjusted.h5ad.gz).

We use single-cell and single-nucleus assays from kidney samples from the Kidney Precision Medicine Project^21^. For this study, we used the single-cell RNA-seq experiments, including 76,020 cells, annotated in 57 major cell types. This data has 36 donors, including 18 control, 5 acute kidney failure, and 13 chronic kidney samples). For this data, we consider those acute kidney failure and chronic kidney samples infected with diabetes. Pre-processed data was obtained from GEO (GSE169285).

The lung cancer single cell atlas is another large data set with 941,504 cells from 165 donors^22^. The samples are formed by 51 normal lung, 18 chronic obstructive pulmonary diseases, 76 lung adenocarcinomas, 13 non-small cell lung carcinomas and 12 squamous cell lung carcinoma. The cells are clustered in 33 clusters/cell types. The atlas is based on distinct studies measured under distinct platforms. We only consider here samples from lung tissue and measured with the 10X genomics platform. Data was obtained from Zenodo (ID 7227571).

We have recently proposed a large study to characterize myocardial infarction^4^. We recovered 115,517 high quality cells, which were clustered in 33 cellular sub-types. We focus here on samples associated with healthy heart tissues (healthy controls and remote zones; n=13) and ischemic zone (IZ; n=7). We exclude one sample due to low quality (less than 1000 genes (on average) per cell). A final dataset is provided in Zenodo (ID 7435911).

The single cell pancreas diabetes data^24^ covers 264,235 mouse pancreatic islet cells (with 13 cell types) from 52 samples. This includes 12 endocrine pancreas disorders, 6 type-1 diabetes mellitus, 12 type-2 diabetes mellitus, and 22 normal samples. We excluded four samples (Embryos E12-E15) due to being outliers by trajectories of methods. The dataset was obtained in GEO (GSE211799). Follicular lymphoma data^23^ collected by Han and colleagues containing 137,147 single cells have been processed to illustrate the various tumor and immune cell populations of Follicular lymphoma. Cells have been clustered into 18 cell types from 23 donors (3 normal and 20 lymphoma samples). The raw data is accessible from EGA (EGAS00001006052).

Finally, we obtained a study with PBMC cells from patients infected with Covid-19^44^. We only considered patients with PMBCs (frozen or fresh cells). This totals to 151 samples classified as either severe infection (n=70), mild infection (n=61), and control (n=20). For these samples, we have 993,171 cells, which were grouped into 10 major cell types. The data was obtained from GEO (GSE158055).

### Pathomics data sets

The VALIGA study is a European cohort with kidney biopsies and accompanying clinical data of patients with IgA nephropathy^25^. We used a pathomics pipeline developed by us^7^ to detect and measure 3 morphometric features of 65,483 tubules and 14 features associated with 24,227 glomeruli structures, which could be detected in 634 biopsies. Patients were classified regarding their glomerular filtration rate (GFR): normal (GFR>60; n=400), reduced (30<GFR<=60; n=177) or low (GFR<=30; n=57). Lower GFR indicates lower kidney function. A second pathomics data set is the Aachen Cohort data, which includes 57 samples with healthy controls and distinct diseases/co-morbidity associated with lower kidney function^7^. We analyzed the histology slides as before^7^, which provided 4,731 glomeruli and 46,999 tubules, each quantified with 3/14 morphometric features. Patients were classified as either being healthy controls (n=17) or diseased (n=40).

We employed a uniform pre-processing pipeline utilizing Seurat^45^ for the normalization and clustering of structures. First, we normalized the data with the function NormalizeData of Seurat, and next ran the ScaleData function. Next, we performed dimension reduction (RunPCA) and kept the 10 and 2 main components for glomeruli and tubules respectively. Next, clusters were found with the Leiden algorithm by using the FindNeighbors and FindClusters functions, respectively. The same pipeline was performed on the kidney IgAN (VALIGA) and Aachen cohort data. As morphometric features are not comparable between glomeruli and tubules, these data are analyzed independently, which results in four distinct data sets. At the patient level, PILOT also allowed the combination of the Wasserstein distance for glomeruli and tubuli, which provided a unique trajectory for each cohort.

### Competing methods

#### Pseudo-bulk

We investigate pseudo-bulk here as a baseline method for sample-level analysis. For single cell data, we sum the gene expression count for all cells in a sample. Then the Poisson distance^26^ between summed counts is employed to compute the sample contrasts. For pathomics data, we calculate the average morphological features per sample and then scale values between 0 and 1. Afterwards, the principal components(PCA) for samples are computed. We detected knees in the variance plots and kept the PCs with the highest variances. We calculated the cosine distance, which was used as input for diffusion maps and Leiden clustering.

#### Proportions

Here, we take the original single cell and pathomics data and calculate the proportion of cell types (clusters) per sample. Next, we calculate the dissimilarity among samples based on their fraction of clusters by cosine distance. Finally, we apply the diffusion map to the distance matrix and get the order of samples.

#### PhEMD

PhEMD required the use of Monocle 2^46^ to perform normalization, dimension reduction, and clustering of cells. The inputs of PhEMD for all data sets are the first 50 principal components of PCA and original pathomic measurements. Among other parameters, PhEMD/Monocle2 requires the definition of a distribution function for expression values. We use as default Negative Binomial distributions to model single cell data sets as advised in the tutorials. For PBMC COVID-19, Diabetic, and Lupus data, we used Gaussianff, due to the fact no cluster was found with the default model. For pathomics data, we used truncated normal distributions due to their best performance. We then used the distance matrices provided by PhEMD as input for a diffusion map analysis as implemented in PILOT.

## Acknowledgements

We thank the European Validation Study of the Oxford Classification of IgAN (VALIGA), funded by the Immunonephrology Working Group of the European Renal Association for access to VALIGA data. This work was funded by grants of the Deutsche Forschungsgemeinschaft (DFG-GE 2811/3) to I.G.C. and the clinical research unit KFO5011 for I.G.C., P.B., R.K. and C.K. and by the Bundesministerium für Bildung und Forschung (BMBF e:Med Consortia Fibromap) to I.G.C. and R.K. PB is supported by the German Research Foundation (DFG, Project IDs 322900939, 454024652, 432698239 445703531), European Research Council (ERC Consolidator Grant No 101001791), and the Federal Ministry of Education and Research (BMBF, STOP-FSGS-01GM2202C).

## Author contributions statement

M.J., M.S., and V.P. developed PILOT. R.B., C.K., D.H., M.G., J.N., N. B., and I.C. performed analysis and interpretation of single cell and pathomics data. M. J., R.K., I.C. conceived the experiments. M.J., M.S. and I.C. wrote the manuscript. All authors reviewed the manuscript.

## Data and code availability

PILOT code, including documentation, tutorials and scripts for replicating experiments, are found in https://github.com/CostaLab/PILOT. Pre-processed R and H5ad objects used as input in benchmarking and case studies are deposited in zenodo, part 1 and zenodo, part 2.

## Group information

Validation Study of the Oxford Classification of IgAN (VALIGA) investigators:

M.L. Russo (MA, PhD, Fondazione Ricerca Molinette, Torino, Italy); S. Troyanov (MD, Division of Nephrology, Department of Medicine, Hopital du Sacre-Coeur de Montreal, Montreal, Quebec, Canada); H.T. Cook (MD, Centre for Complement and Inflammation Research, Department of Medicine, Imperial College, London, England); I. Roberts (MD, Department of Cellular Pathology, Oxford University Hospitals NHS Foundation Trust, John Radcliffe Hospital, Oxford, United Kingdom); V. Tesar, (MD, Department of Nephrology, 1st Faculty of Medicine and General University Hospital, Charles University, Prague, Czech Republic); D. Maixnerova (MD, Department of Nephrology, 1st Faculty of Medicine and General University Hospital, Charles University, Prague, Czech Republic); S. Lundberg (MD, Nephrology Unit, Department of Clinical Sciences, Karolinska Institute, Stockholm, Sweden); L. Gesualdo (MD, Department of Nephrology, Emergency and Organ Transplantation, University of Bari “Aldo Moro,” Foggia-Bari, Italy); F. Emma (MD, Division of Nephrology, Department of Pediatric Subspecialties, Bambino Gesù Children’s Hospital IRCCS, Rome, Italy); F. Diomedi (MD, Division of Nephrology, Department of Pediatric Subspecialties, Bambino Gesù Children’s Hospital IRCCS, Rome, Italy); G. Beltrame (MD, Nephrology and Dialysis Unit, San Giovanni Bosco Hospital, and University of Turin,Turin, Italy); C. Rollino (MD, Nephrology and Dialysis Unit, San Giovanni Bosco Hospital, and University of Turin,Turin, Italy); A. Amore (MD, Nephrology Unit, Regina Margherita Children’s Hospital,Turin, Italy); R. Camilla (MD Nephrology Unit, Regina Margherita Children’s Hospital, Turin, Italy); L. Peruzzi (MD, Nephrology Unit, Regina Margherita Children’s Hospital, Turin, Italy); M. Praga (MD, Nephrology Unit, Hospital 12 de Octubre,Madrid, Spain); S. Feriozzi (MD, Nephrology Unit, Belcolle Hospital, Viterbo, Italy), R. Polci, (MD, Nephrology Unit, Belcolle Hospital,Viterbo, Italy); G. Segoloni, (MD, Division of Nephrology Dialysis and Transplantation, Department of Medical Sciences, Città della Salute e della Scienza Hospital and University of Turin, Turin, Italy); L.Colla (MD, Division of Nephrology Dialysis and Transplantation, Department of Medical Sciences, Città della Salute e della Scienza Hospital and University of Turin,Turin, Italy); A. Pani (MD, Nephrology Unit, G. Brotzu Hospital, Cagliari, Italy); D. Piras (MD, Nephrology Unit, G. Brotzu Hospital, Cagliari, Italy), A. Angioi (MD, Nephrology Unit, G. Brotzu Hospital, Cagliari, Italy); G. Cancarini, (MD, Nephrology Unit, Spedali Civili University Hospital, Brescia, Italy); S. Ravera (MD, Nephrology Unit, Spedali Civili University Hospital, Brescia, Italy); M. Durlik (MD, Department of Transplantation Medicine, Nephrology, and Internal Medicine, Medical University ofWarsaw,Warsaw, Poland); E. Moggia (Nephrology Unit, Santa Croce Hospital, Cuneo, Italy); J. Ballarin (MD, Department of Nephrology, Fundacion Puigvert, Barcelona, Spain); S. Di Giulio (MD, Nephrology Unit, San Camillo Forlanini Hospital, Rome, Italy); F. Pugliese (MD, Department of Nephrology, Policlinico Umberto I University Hospital, Rome, Italy); I. Serriello (MD, Department of Nephrology, Policlinico Umberto I University Hospital, Rome, Italy); Y. Caliskan (MD, Division of Nephrology, Department of Internal Medicine, Istanbul Faculty of Medicine, Istanbul University, Istanbul, Turkey); M. Sever (MD, Division of Nephrology, Department of Internal Medicine, Istanbul Faculty of Medicine, Istanbul University, Istanbul, Turkey); I. Kilicaslan (MD, Department of Pathology, Istanbul Faculty of Medicine, Istanbul University, Istanbul, Turkey); F. Locatelli (MD, Department of Nephrology and Dialysis, Alessandro Manzoni Hospital, ASST Lecco, Italy); L. Del Vecchio (MD, Department of Nephrology and Dialysis, Alessandro Manzoni Hospital, ASST Lecco, Italy); J.F.M.Wetzels (MD, Departments of Nephrology, Radboud University Medical Center, Nijmegen, the Netherlands); H. Peters (MD, Departments of Nephrology, Radboud University Medical Center, Nijmegen, the Netherlands); U. Berg (MD, Division of Pediatrics, Department of Clinical Science, Intervention and Technology, Huddinge, Sweden); F. Carvalho (MD, Nephrology Unit, Hospital de Curry Cabral, Lisbon, Portugal); A.C. da Costa Ferreira (MD, Nephrology Unit, Hospital de Curry Cabral, Lisbon, Portugal); M. Maggio (MD, Nephrology Unit, Hospital Maggiore di Lodi, Lodi, Italy); A. Wiecek (MD, Department Nephrology, Endocrinology and Metabolic Diseases, Silesian University of Medicine, Katowice, Poland); M. Ots-Rosenberg(MD, Nephrology Unit, Tartu University Clinics, Tartu,Estonia); R. Magistroni (MD, Department of Nephrology, Policlinic of Modena and Reggio Emilia; Modena, Italy); R. Topaloglu (MD, Department of Pediatric Nephrology and Rheumatology, Hacettepe University, Ankara, Turkey); Y. Bilginer (MD, Department of Pediatric Nephrology and Rheumatology, Hacettepe University, Ankara, Turkey); M. D’Amico (MD, Nephrology Unit, S. Anna Hospital, Como, Italy); M. Stangou (MD, Department of Nephrology, Hippokration General Hospital, Aristotle University of Thessaloniki, Thessaloniki, Greece); F. Giacchino (MD, Nephrology Unit, Ivrea Hospital, Ivrea, Italy); D. Goumenos (MD Department of Nephrology, University Hospital of Patras, Patras, Greece); P. Kalliakmani (MD Department of Nephrology, University Hospital of Patras, Patras, Greece); M. Papasotiriou (MD Department of Nephrology, University Hospital of Patras, Patras, Greece); K. Galesic (MD, Department of Nephrology, University Hospital Dubrava, Zagreb, Croatia); C. Geddes (MD, Renal Unit,Western Infirmary Glasgow, Glasgow, United Kingdom); K. Siamopoulos (MD, Nephrology Unit, Medical School University of Ioanina, Ioannina, Greece); O. Balafa (MD, Nephrology Unit,Medical School University of Ioanina, Ioannina, Greece); M. Galliani (MD, Nephrology Unit, S.Pertini Hospital, Rome, Italy); P. Stratta (MD, Department of Nephrology, Maggiore della Carità Hospital, Piemonte Orientale University, Novara, Italy); M. Quaglia (MD, Department of Nephrology, Maggiore della Carità Hospital, Piemonte Orientale University, Novara, Italy); R. Bergia (MD, Nephrology Unit, Degli Infermi Hospital, Biella,Italy); R. Cravero (MD, Nephrology Unit, Degli Infermi Hospital, Biella, Italy); M. Salvadori, (MD, Department of Nephrology, Careggi Hospital, Florence, Italy); L. Cirami (MD, Department of Nephrology, Careggi Hospital, Florence, Italy); B. Fellstrom (MD, Renal Department, University of Uppsala, Uppsala, Sweden); H. Kloster Smerud (MD, Renal Department, University of Uppsala, Uppsala, Sweden); F. Ferrario (MD, Nephropathology Unit, San Gerardo Hospital, Monza, Italy); T. Stellato (MD, Nephropathology Unit, San Gerardo Hospital, Monza, Italy); J. Egido (MD, Department of Nephrology, Fundacion Jimenez Diaz, Madrid, Spain); C. Martin (MD, Department of Nephrology, Fundacion Jimenez Diaz, Madrid, Spain); J. Floege (MD, Nephrology and Immunology, Medizinische Klinik II, University of Aachen, Aachen, Germany); F. Eitner (MD, Nephrology and Immunology, Medizinische Klinik II, University of Aachen, Aachen, Germany); A. Lupo (MD, Department of Nephrology, University of Verona, Verona, Italy); P. Bernich (MD, Department of Nephrology, University of Verona, Verona, Italy); P. Menè (Department of Nephrology,S. Andrea Hospital, Rome, Italy); M. Morosetti (Nephrology Unit, Grassi Hospital, Ostia, Italy); C. van Kooten, (MD, Department of Nephrology, Leiden University Medical Centre, Leiden, The Netherlands); T. Rabelink (MD, Department of Nephrology, Leiden University Medical Centre, Leiden, The Netherlands); M.E.J. Reinders (MD, Department of Nephrology, Leiden University Medical Centre, Leiden, The Netherlands); J.M. Boria Grinyo (Department of Nephrology, Hospital Bellvitge, Barcelona, Spain); S. Cusinato (MD, Nephrology Unit, Borgomanero Hospital, Borgomanero, Italy); L. Benozzi (MD, Nephrology Unit, Borgomanero Hospital, Borgomanero, Italy); S. Savoldi, (MD, Nephrology Unit, Civile Hospital, Ciriè, Italy); C. Licata (MD, Nephrology Unit, Civile Hospital, Ciriè, Italy); M. Mizerska-Wasiak (MD, Department of Pediatrics, Medical University ofWarsaw,Warsaw, Poland); G. Martina (MD, Nephrology Unit, Chivasso Hospital, Chivasso, Italy); A. Messuerotti (MD, Nephrology Unit, Chivasso Hospital, Chivasso, Italy); A. Dal Canton (MD, Nephrology Unit, S. Matteo Hospital, Pavia, Italy); C. Esposito (MD, Nephrology Unit, Maugeri Foundation, Pavia, Italy); C. Migotto (MD, Nephrology Unit, Maugeri Foundation, Pavia, Italy); G. Triolo MD, Nephrology Unit CTO, Turin, Italy); F.Mariano (MD, Nephrology Unit CTO, Turin, Italy); C. Pozzi (MD, Nephrology Unit, Bassini Hospital, Cinisello Balsamo, Italy); R. Boero (MD, Nephrology Unit, Martini Hospital, Turin, Italy);

VALIGA pathology investigators: S. Bellur (MD, Department of Cellular Pathology, Oxford University Hospitals NHS Foundation Trust, John Radcliffe Hospital, Oxford, United Kingdom); G.Mazzucco (MD, Pathology Department, University of Turin, Turin, Italy); C. Giannakakis (MD, Pathology Department, La Sapienza University, Rome, Italy); E. Honsova (MD, Department of Clinical and Transplant Pathology, Institute for Clinical and Experimental Medicine, Prague, Czech Republic); B. Sundelin (MD Department of Pathology and Cytology, Karolinska University Hospital, Karolinska Institute, Stockholm, Sweden); A.M. Di Palma (Nephrology Unit, Aldo Moro University, Foggia-Bari, Italy); F. Ferrario (MD, Nephropathology Unit, San Gerardo Hospital, Monza, Italy); E. Gutiérrez (MD, Renal, Vascular and Diabetes Research Laboratory, Fundación Instituto de Investigaciones Sanitarias-Fundación Jiménez Díaz, Universidad Autónoma de Madrid, Madrid, Spain); A.M. Asunis (MD, Department of Pathology, Brotzu Hospital, Cagliari, Italy); J. Barratt (MD, The JohnWalls Renal Unit, Leicester General Hospital, Leicester, United Kingdom); R. Tardanico (MD, Department of Pathology, Spedali Civili Hospital, University of Brescia, Brescia, Italy); A. Perkowska-Ptasinska (MD, Department of Transplantation Medicine, Nephrology and Internal Medicine, Medical University of Warsaw, Warsaw, Poland); J. Arce Terroba (MD, Pathology Department, Fundació Puigvert, Barcelona, Spain); M. Fortunato (MD, Pathology Department, S. Croce Hospital, Cuneo, Italy); A. Pantzaki (MD, Department of Pathology, Hippokration Hospital, Thessaloniki, Greece); Y. Ozluk (MD, Department of Pathology, Istanbul University, Istanbul Faculty of Medicine, Istanbul, Turkey); E. Steenbergen (MD, Radboud University Medical Center, Department of Pathology, Nijmegen, The Netherlands); M. Soderberg (MD, Department of Pathology, Drug Safety and Metabolism, Huddinge, Sweden); Z. Riispere (MD, Department of Pathology, University of Tartu, Tartu, Estonia); L. Furci (MD, Pathology Department, University of Modena, Italy); D. Orhan (MD, Department of Pediatrics, Division of Rheumatology, Hacettepe University Faculty of Medicine, Ankara, Turkey); D. Kipgen (MD,Pathology Department, Queen Elizabeth University Hospital, Glasgow, United Kingdom); D. Casartelli (Pathology Department, Manzoni Hospital, Lecco, Italy); D. Galesic Ljubanovic (MD, Nephrology Department, University Hospital, Zagreb, Croatia; Zagreb, Croatia); H Gakiopoulou (MD, Department of Pathology, National and Kapodistrian University of Athens, Athens, Greece); E. Bertoni (MD, Nephrology Department, Careggi Hospital, Florence, Italy); P. Cannata Ortiz (MD, Pathology Department, IIS-Fundacion Jimenez Diaz UAM, Madrid, Spain); H. KarkoszkaMD, (Nephrology, Endocrinology and Metabolic Diseases, Medical University of Silesia, Katowice, Katowice, Poland); H.J. Groene (MD, Cellular and Molecular Pathology, German Cancer Research Center, Heidelberg, Germany); A. Stoppacciaro (MD, Surgical Pathology Units, Department of Clinical and Molecular Medicine, Ospedale Sant’Andrea, Sapienza University of Rome, Rome, Italy); I. Bajema (MD, Department of Pathology, Leiden University Medical Center, Leiden, The Netherlands); J. Bruijn (MD, Department of Pathology, Leiden University Medical Center, Leiden, The Netherlands); X. Fulladosa Oliveras (MD, Nephrology Unit, Bellvitge University Hospital, Hospitalet de Llobregat, Barcelona, Spain); J. Maldyk (MD, Division of Pathomorphology, Children’s Clinical Hospital, Medical University ofWarsaw,Warsaw, Poland); and E. Ioachim (MD, Department of Pathology, Medical School, University of Ioannina, Ioannina, Greece); the Oxford derivation and North American validation investigators: Bavbek N (MD, Department of Pathology, Vanderbilt University, Nashville, Tennessee); Cook T (MD, Imperial College, London, England), Troyanov S (MD, Division of Nephrology, Department of Medicine, Hopital du Sacre-Coeur de Montreal, Montreal, Quebec, Canada); Alpers C (MD, Department of Pathology, University of Washington Medical Center, Seattle,Washington), Amore A (MD, Nephrology, Dialysis and Transplantation Unit, Regina Margherita Children’s Hospital, University of Turin, Turin, Italy), Barratt J (MD, The John Walls Renal Unit, Leicester General Hospital, Leicester, England); Berthoux F (MD, Department of Nephrology, Dialysis, and Renal Transplantation, Hôpital Nord, CHU de Saint-Etienne, Saint-Etienne, France); Bonsib S (MD, Department of Pathology, LSU Health Sciences Center, Shreveport, Los Angeles); Bruijn J (MD, Department of Pathology, Leiden University Medical Center, Leiden, The Netherlands); D’Agati V (MD, Department of Pathology, Columbia University College of Physicians Surgeons, New York, New York); D’Amico G (MD, Fondazione D’Amico per la Ricerca sulle Malattie Renali, Milan, Italy); Emancipator S (MD, Department of Pathology, Case Western Reserve University, Cleveland, Ohio); Emmal F (MD, Division of Nephrology and Dialysis, Department of Nephrology and Urology, Bambino Gesù Children’s Hospital and Research Institute, Piazza S Onofrio, Rome, Italy); Ferrario F (MD, Renal Immunopathology Center, San Carlo Borromeo Hospital, Milan, Italy); Fervenza F (MD PhD, Division of Nephrology and Hypertension, Mayo Clinic, Rochester); Florquin S (MD, Department of Pathology, Academic Medical Center, University of Amsterdam, Amsterdam, The Netherlands); Fogo A (MD, Department of Pathology, Vanderbilt University, Nashville, Tennessee); Geddes C (MD, The Renal Unit, Western Infirmary, Glasgow, Scotland); Groene H (MD, Department of Cellular and Molecular Pathology, German Cancer Research Center, Heidelberg, Germany); Haas M(MD, Department of Pathology and Laboratory Medicine, Cedars-Sinai Medical Center, Los Angeles, California); Hill P (MD, St Vincent’s Hospital, Melbourne, Australia); Hogg R (MD, Scott and White Medical Center, Temple, Texas (retired)); Hsu S (MD, Division of Nephrology, Hypertension and Renal Transplantation, College of Medicine, University of Florida, Gainesville, Florida); Hunley T (MD, Department of Pathology, Vanderbilt University, Nashville, Tennessee); Hladunewich (MD, Division of Nephrology, Sunnybrook Health Science Center, University of Toronto, Ontario, Canada M); Jennette C (MD, Department of Pathology and Laboratory Medicine, University of North Carolina, Chapel Hill, North Carolina); Joh K (MD, Division of Immunopathology, Clinical Research Center Chiba, East National Hospital, Chiba, Japan); Julian B (MD, Department of Medicine, University of Alabama at Birmingham, Birmingham, Alabama); Kawamura T (MD, Division of Nephrology and Hypertension, Jikei University School of Medicine, Tokyo, Japan); Lai F (MD, The Chinese University of Hong Kong, Hong Kong); Leung C (MD, Department of Medicine, Prince of Wales Hospital, Chinese University of Hong Kong, Hong Kong); Li L (MD, Research Institute of Nephrology, Jinling Hospital, Nanjing University School of Medicine, Nanjing, China); Li P (MD, Department of Medicine, Prince of Wales Hospital, Chinese University of Hong Kong, Hong Kong); Liu Z (MD, Research Institute of Nephrology, Jinling Hospital, Nanjing University School of Medicine, Nanjing, China); Massat A (MD, Division of Nephrology and Hypertension, Mayo Clinic, Rochester, Minnesota); Mackinnon B (MD, The Renal Unit,Western Infirmary, Glasgow, Scotland); Mezzano S (MD, Departamento de Nefrología, Escuela de Medicina, Universidad Austral, Valdivia, Chile); Schena F (MD, Renal, Dialysis and Transplant Unit, Policlinico, Bari, Italy); Tomino Y (MD, Division of Nephrology, Department of Internal Medicine, Juntendo University School of Medicine, Tokyo, Japan); Walker P (MD, Nephropathology Associates, Little Rock, Arkansas);Wang H (MD, Renal Division of Peking University First Hospital, Peking University Institute of Nephrology, Beijing, China (deceased)); Weening J (MD, Erasmus Medical Center, Rotterdam, The Netherlands); and Yoshikawa N (MD, Department of Pediatrics, Wakayama Medical University, Wakayama City, Japan); the International investigators: Cai-Hong Zeng (MD, Nanjing University School of Medicine, Nanjing, China); Sufang Shi (MD, Peking University Institute of Nephrology, Beijing, China); C.Nogi (MD, Juntendo University, Faculty of Medicine, Tokyo, Japan); H.Suzuki (MD, Juntendo University, Faculty of Medicine, Tokyo, Japan); K. Koike (MD, Division of Nephrology and Hypertension, Department of Internal Medicine, Jikei University School of Medicine, Tokyo, Japan); K. Hirano (MD, Division of Nephrology and Hypertension, Department of Internal Medicine, Jikei University School of Medicine, Tokyo, Japan); T. Kawamura (MD, Division of Nephrology and Hypertension, Department of Internal Medicine, Jikei University School of Medicine, Tokyo, Japan); T. Yokoo (MD, Division of Nephrology and Hypertension, Department of Internal Medicine, Jikei UniversitySchool of Medicine, Tokyo, Japan); M. Hanai (MD, Division of Nephrology, Department of Medicine, Kurume University School of Medicine, Fukuoka, Japan); K. Fukami (MD, Division of Nephrology, Department of Medicine, Kurume University School of Medicine, Fukuoka„ Japan); K. Takahashi (MD, Department of Nephrology, Fujita Health University School of Medicne, Aichi, Japan); Y. Yuzawa (MD, Department of Nephrology, Fujita Health University School of Medicine, Aichi, Japan); M. Niwa (MD, Department of Nephrology, Nagoya University Graduate School of Medicine, Aichi, Japan); Y. Yasuda (MD, Department of Nephrology, Nagoya University Graduate School of Medicine, Aichi, Japan); S. Maruyama (MD, Department of Nephrology, Nagoya University Graduate School of Medicine, Aichi, Japan); D. Ichikawa (MD, Division of Nephrology and Hypertension, Department of Internal Medicine, St Marianna University School of Medicine, Kanagawa, Japan); T. Suzuki (MD, Division of Nephrology and Hypertension, Department of Internal Medicine, St Marianna University School of Medicine, Kanagawa, Japan); S. Shirai (MD, Division of Nephrology and Hypertension, Department of Internal Medicine, St Marianna University School of Medicine, Kanagawa, Japan); A. Fukuda (MD, First Department of Internal Medicine, Faculty of Medicine, University of Miyazaki, Miyazaki, Japan); S. Fujimoto (MD, Department of Hemovascular Medicine and Artificial Organs, Faculty of Medicine, University of Miyazaki, Miyazaki, Japan); H. Trimarchi (MD, Division of Nephrology, Hospital Britanico, Buenos Aires, Argentina).

## Footnotes

1 This computing time of PhEMD also includes clustering of single cells, as this is a required step in PhEMD framework

2 For simplicity, we focus here on single cell data, but the same formalism applies to pathomics data

## Notes

https://github.com/CostaLab/PILOT

### Competing Interest Statement

The authors have declared no competing interest.

### Summary of Updates

We have expanded the number of data sets, expanded the benchmark with additional baseline methods and improved the text.

