## Supplementary material for "Detection of PatIent-Level distances from single cell genomics and pathomics data with Optimal Transport (PILOT)": Sup. Table and Figures

### Supplementary Tables and Figures

**Table S1.** Benchmarking of clustering results (ARI and Silhouette Scores), and area under the PR curve (AUPR). Results in bold indicate the best value per method.

| **Methods** | **Pseudobulk** | | | **Phemd** | | | **Proportions** | | | **PILOT** | | |
|---|---|---|---|---|---|---|---|---|---|---|---|---|
| Datasets/Metrics | ARI | Sil | AUCPR | ARI | Sil | AUCPR | ARI | Sil | AUCPR | ARI | Sil | AUCPR |
| PDAC | **1.00** | **0.73** | **1.00** | 0.51 | 0.04 | 0.87 | 0.51 | 0.56 | **1.00** | **1.00** | 0.59 | **1.00** |
| Lupus PBMC | 0.57 | 0.34 | **0.92** | 0.51 | 0.02 | 0.58 | **0.70** | **0.39** | 0.91 | 0.61 | 0.28 | 0.89 |
| Kidney IgAN G | 0.50 | 0.01 | 0.51 | 0.50 | -0.04 | 0.34 | 0.51 | 0.00 | 0.55 | **0.55** | **0.07** | **0.60** |
| Myoc. Infarc. | **1.00** | **0.86** | **1.00** | 0.47 | -0.06 | 0.26 | 0.81 | 0.70 | 0.98 | 0.90 | 0.73 | 0.98 |
| Kidney AC T | 0.57 | 0.21 | 0.54 | 0.51 | 0.00 | 0.76 | 0.53 | 0.25 | 0.87 | **0.68** | **0.30** | **0.93** |
| Kidney AC G | 0.51 | 0.25 | 0.59 | 0.51 | -0.01 | 0.66 | 0.57 | 0.25 | **0.96** | **0.64** | **0.36** | **0.96** |
| COVID-19 | 0.51 | -0.25 | 0.92 | 0.51 | **-0.06** | 0.88 | 0.52 | -0.14 | **0.95** | **0.55** | -0.14 | 0.94 |
| Kidney | 0.71 | 0.16 | 0.77 | **0.76** | **0.44** | **1.00** | 0.70 | 0.24 | 0.92 | 0.62 | 0.11 | 0.81 |
| Kidney IgAN T | 0.48 | 0.07 | 0.54 | 0.49 | -0.03 | 0.48 | 0.56 | 0.00 | 0.54 | **0.58** | **0.10** | **0.59** |
| Lung | **0.70** | -0.23 | 0.87 | 0.29 | -0.02 | 0.85 | 0.67 | -0.07 | **0.90** | 0.67 | **-0.01** | **0.90** |
| Diabetes | **0.77** | **0.18** | 0.70 | 0.62 | -0.13 | 0.51 | 0.64 | -0.03 | 0.50 | 0.67 | -0.01 | **0.71** |
| Foll. lym. | 0.50 | 0.29 | 0.90 | 0.52 | 0.09 | 0.97 | 0.48 | **0.95** | **1.00** | **0.76** | 0.60 | **1.00** |

**Table S2.** Log likelihood test and hazard ratios (HR) of cox regression for disease progression and morphometric variables

| **Features** | **p_value** | **HR** | **HR_lower_95** | **HR_higher_95** |
|---|---|---|---|---|
| Disease_Progression | 2.38E-11 | 14.22285964 | 6.133761376 | 32.97972059 |
| Tuft-Area-Fraction | 6.55E-06 | 0.006096 | 0.000754147 | 0.049275825 |
| Age | 0.000161201 | 1.026118273 | 1.012515212 | 1.03990409 |
| Tuft Elongation | 0.035174436 | 0.995855111 | 0.991024645 | 1.000709122 |
| Tuft Area | 0.059838632 | 0.99999991 | 0.999999796 | 1.000000024 |
| Glomerular Area | 0.084688986 | 0.99999995 | 0.999999883 | 1.000000017 |
| Tuft Circularity | 0.120647357 | 0.997847514 | 0.99474223 | 1.000962492 |
| Tuft Eccentricity | 0.193320223 | 0.99911608 | 0.99765839 | 1.0005759 |
| Bowman Area | 0.285387443 | 0.999999915 | 0.999999741 | 1.000000088 |
| Glomerular Circularity | 0.292065233 | 0.998989099 | 0.996929139 | 1.001053316 |
| Glomerular Diameter | 0.295112885 | 0.999996112 | 0.999988267 | 1.000003957 |
| Tubular Distance | 0.334379046 | 0.999806135 | 0.999357973 | 1.000254498 |
| Tubular Area | 0.361417305 | 1.000000128 | 0.999999865 | 1.000000392 |
| Tuft Solidity | 0.468203891 | 0.999443691 | 0.997818499 | 1.001071531 |
| Glomerular Distance | 0.55738282 | 1.000000526 | 0.999998905 | 1.000002148 |
| Glomerular Solidity | 0.594992091 | 0.999714882 | 0.998620471 | 1.000810492 |
| Glomerular Eccentricity | 0.721397871 | 1.000225975 | 0.999008779 | 1.001444655 |
| Glomerular Elongation | 0.944083375 | 0.999862207 | 0.995995708 | 1.003743715 |
| Tubular Diameter | 0.954027614 | 0.999999227 | 0.999972858 | 1.000025597 |

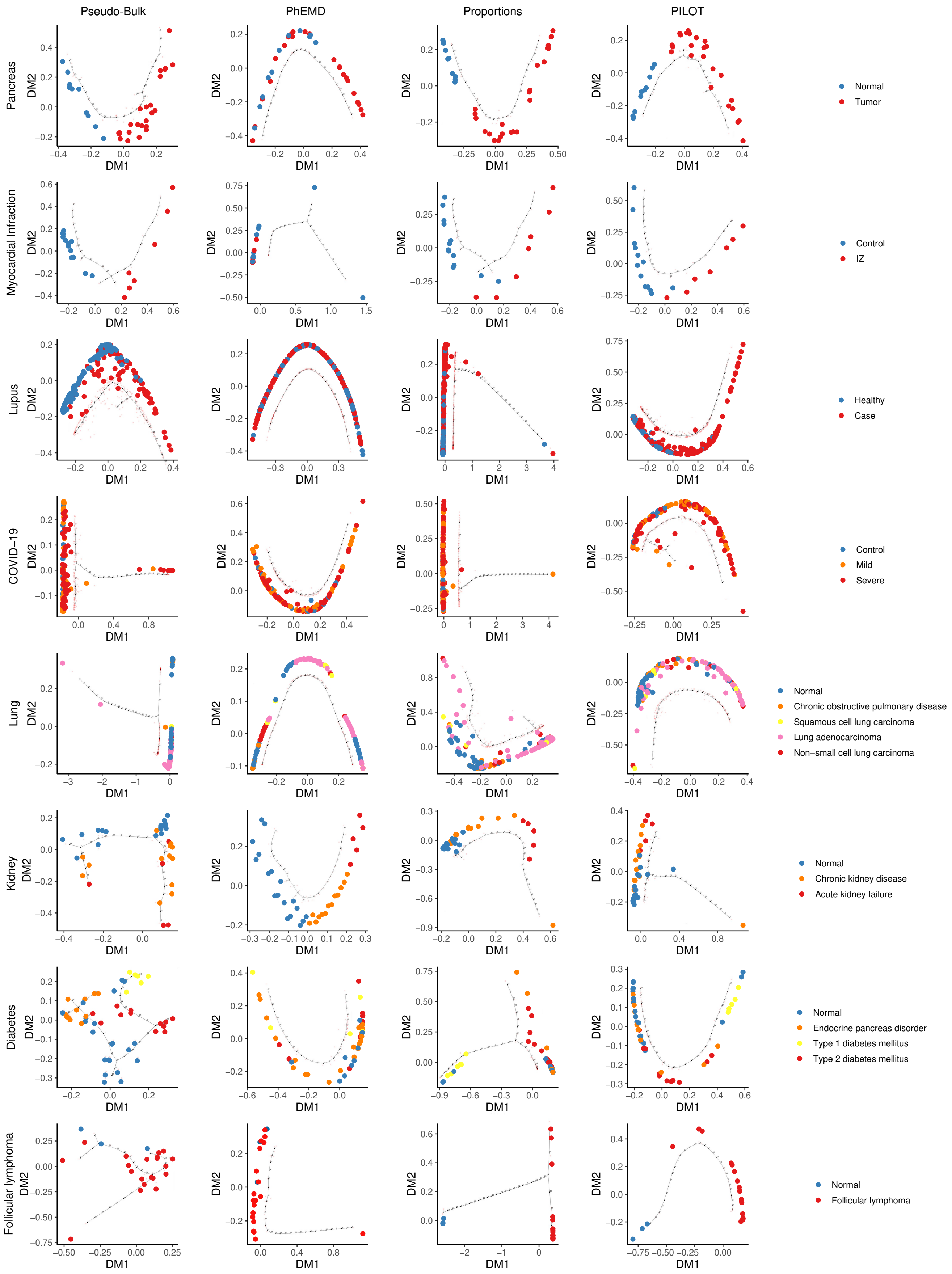


**Figure S1.** Diffusion maps of single cell data sets (rows) for Pseudo Bulk, PhEMD, Proportions, and PILOT (columns). Doted lines represent backbones of the estimated trajectories.

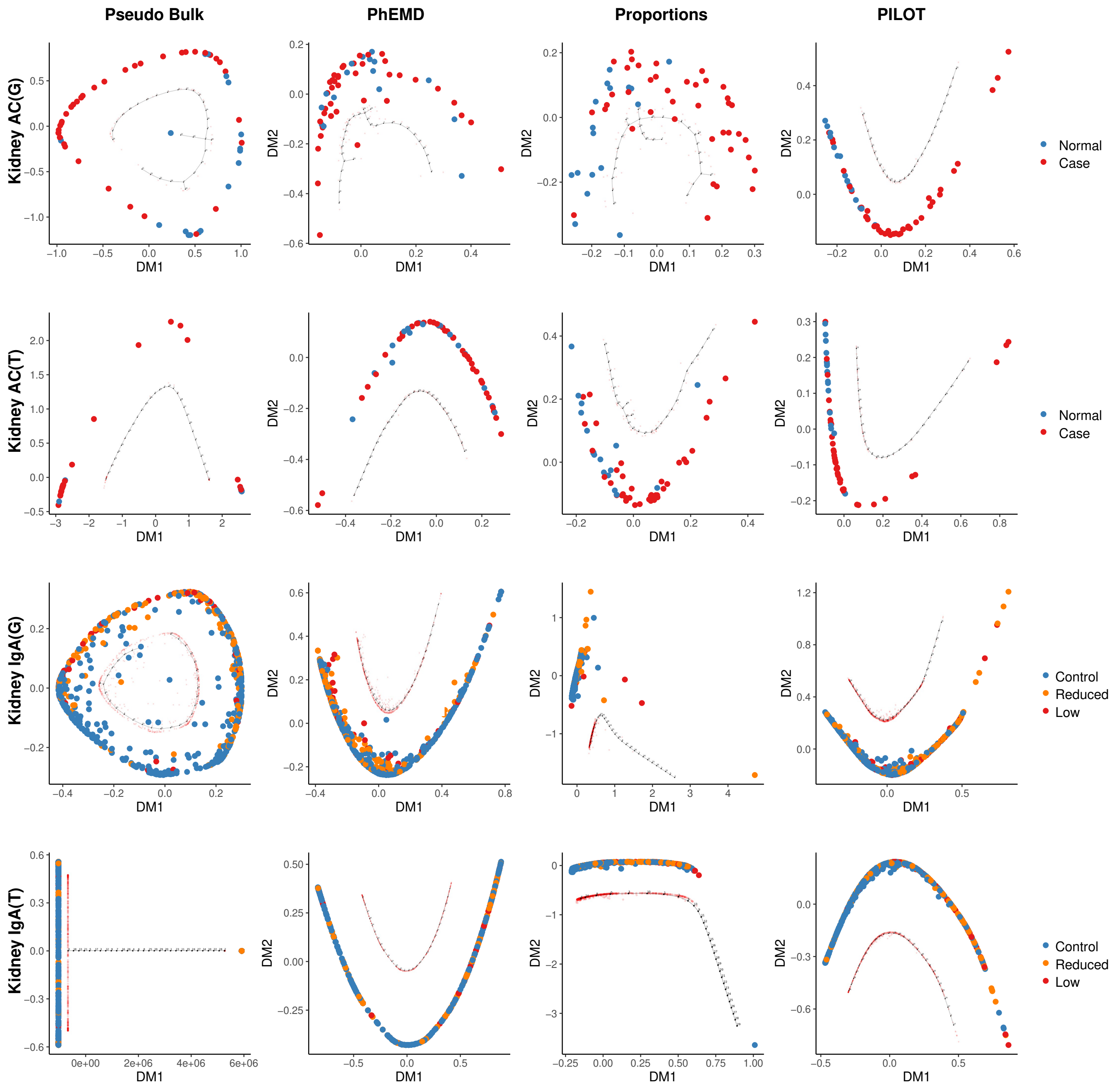


**Figure S2.** Diffusion maps of pathomics data sets (rows) for Pseudo Bulk, PhEMD, Proportions, and PILOT (columns). Doted lines represent backbones of the estimated trajectories.

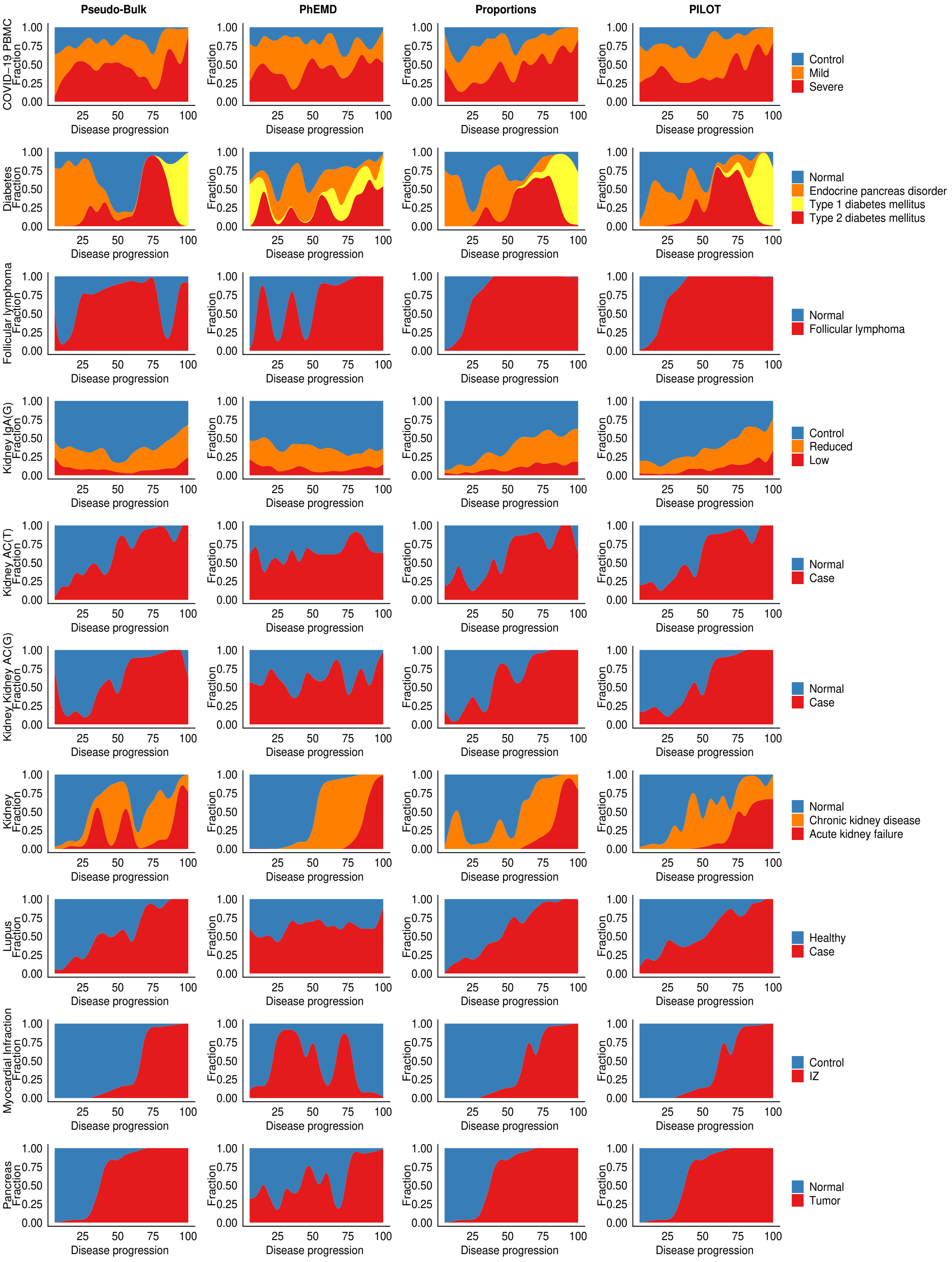


**Figure S3.** Fraction samples (y-axis) for distinct sample labels over the pseudo-time (x-axis) for disease progression trajectories.

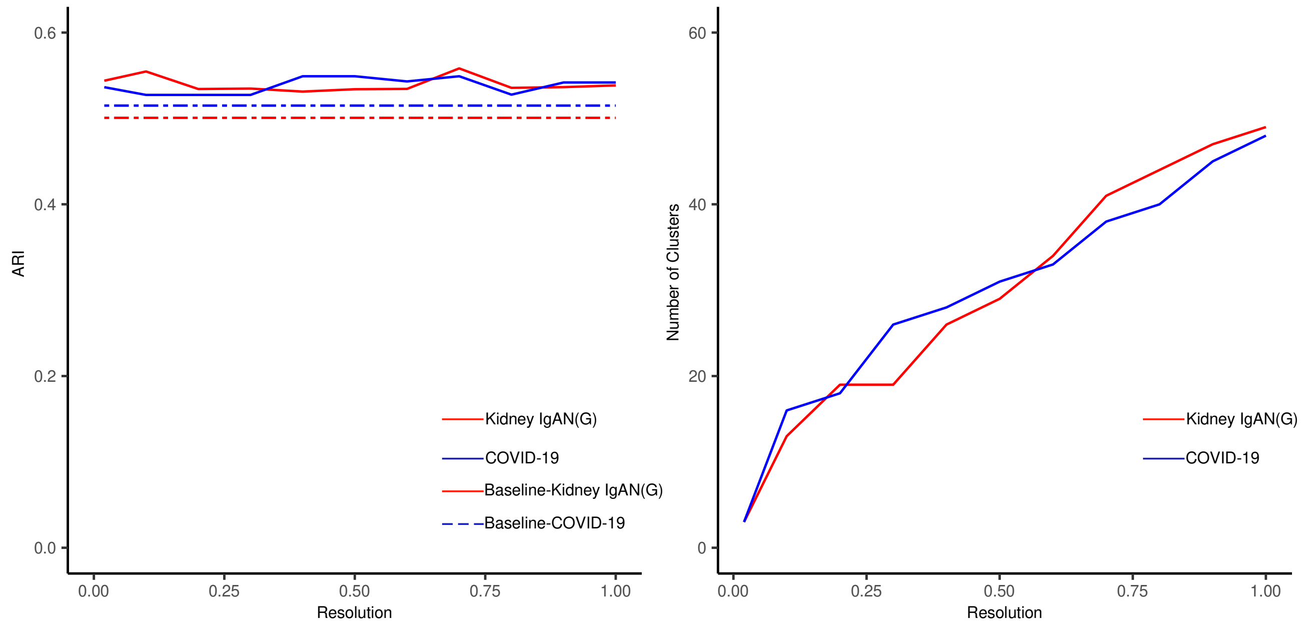


**Figure S4.** A - Clustering performance (measured by ARI) for PILOT (solid lines) and pseudo-bulk (traced lines) for distinct cluster resolution. We observe that for both Kidney IgAN and COVID-19, the number of clusters did not impacted the results from PILOT. B - Number of clusters for distinct Leiden algorithm resolution parameters.

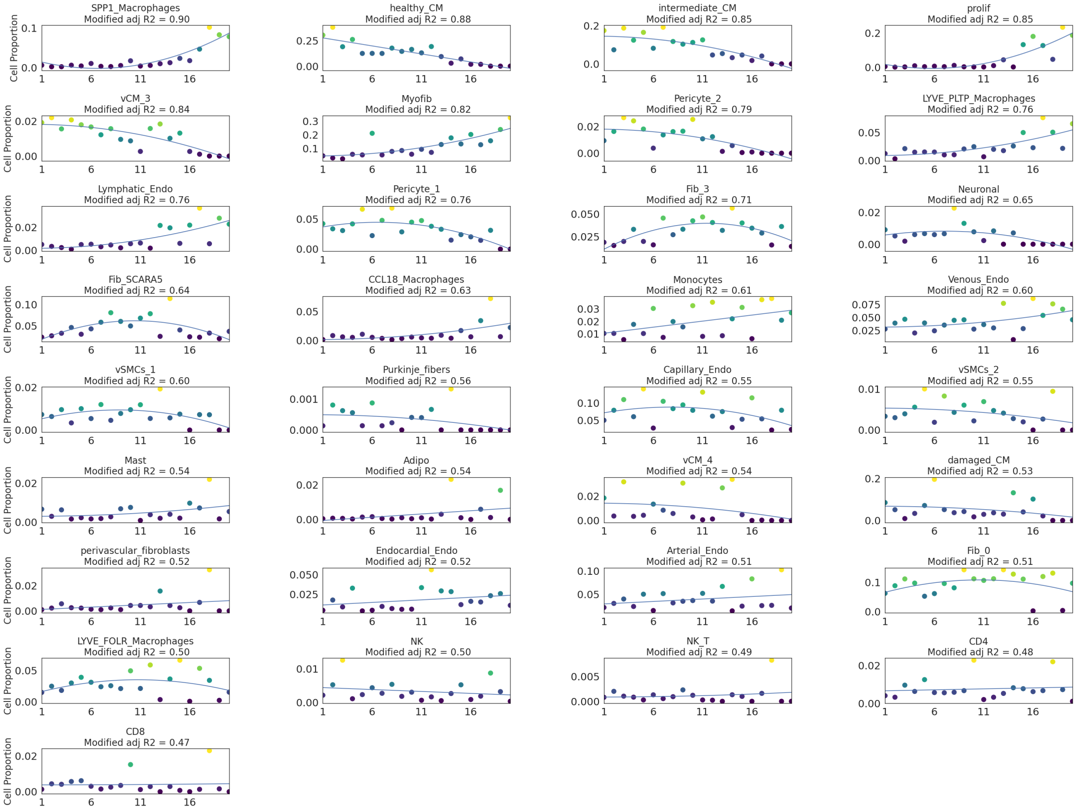


**Figure S5.** Cell type frequency (y-axis) vs. PILOT disease progression (x-axis) for Myocardial Infarction scRNA-seq data.

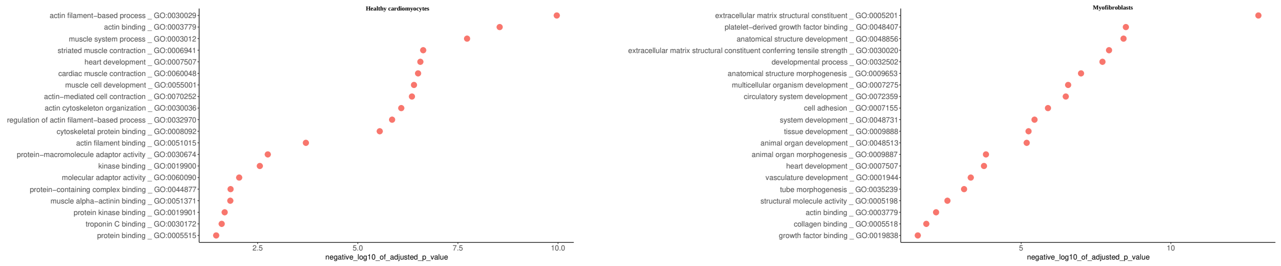


**Figure S6.** Top 20 terms of the enrichment analysis by using selected genes for Healthy cardiomyocytes and myofibroblasts (based on the first 100 genes with Larger FC and $p$-value $< 0.001$). The analysis was performed with g:profiler `https://biit.cs.ut.ee/gprofiler/gost`.

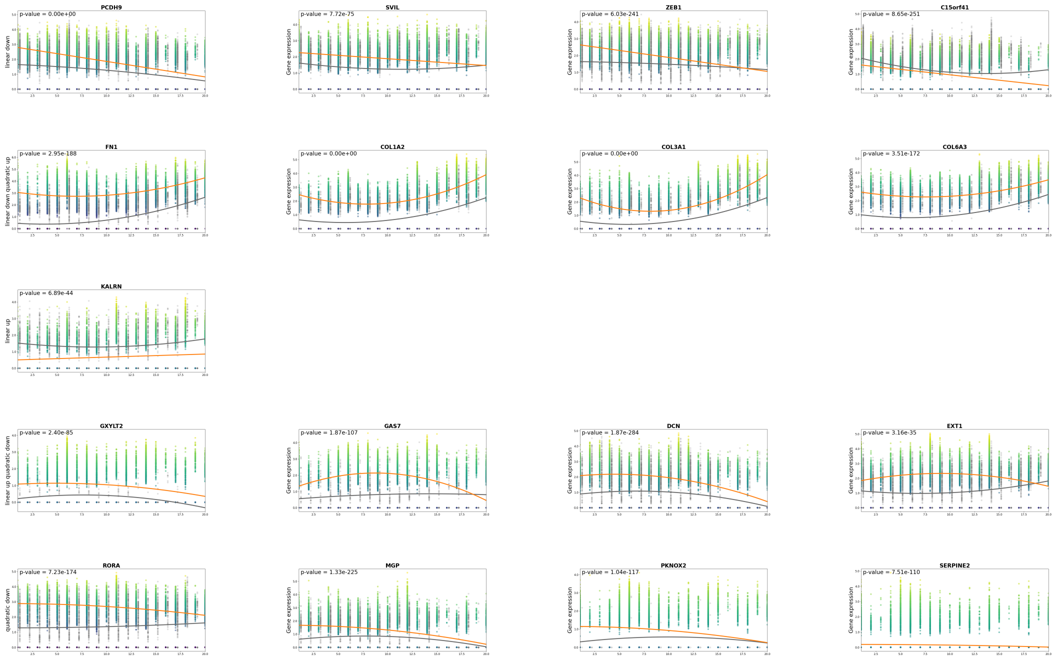


**Figure S7.** Top genes associated (ranked by FC) from Myofibroblasts of the Myocardial infarction scRNA-seq. Every line corresponds to distinct models (linear, quadratic, linear-quadratic) and patterns (up vs. down). Only significant genes ($p$-value $> 0.01$) are shown.

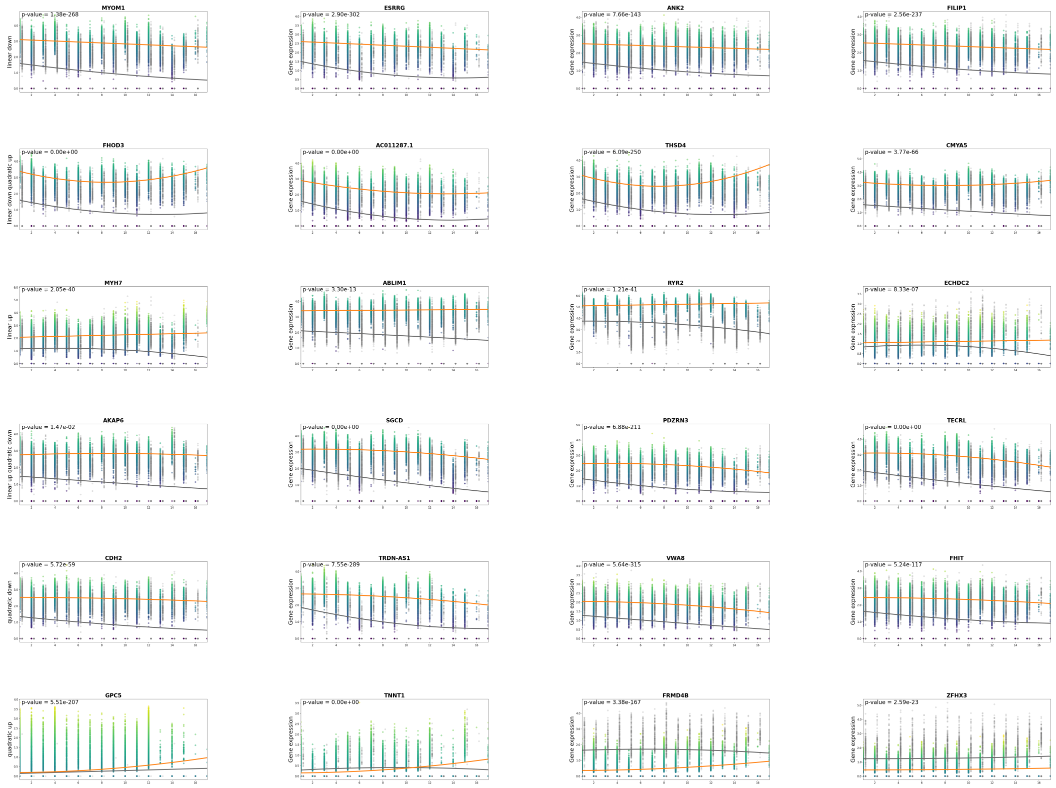


**Figure S8.** Top genes associated (ranked by FC) from Healthy cardiomiocytes of the myocardial infarction scRNA-seq. Every line corresponds to distinct models (linear, quadratic, linear-quadratic) and patterns (up vs. down). Only significant genes ($p$-value > 0.01) are shown.

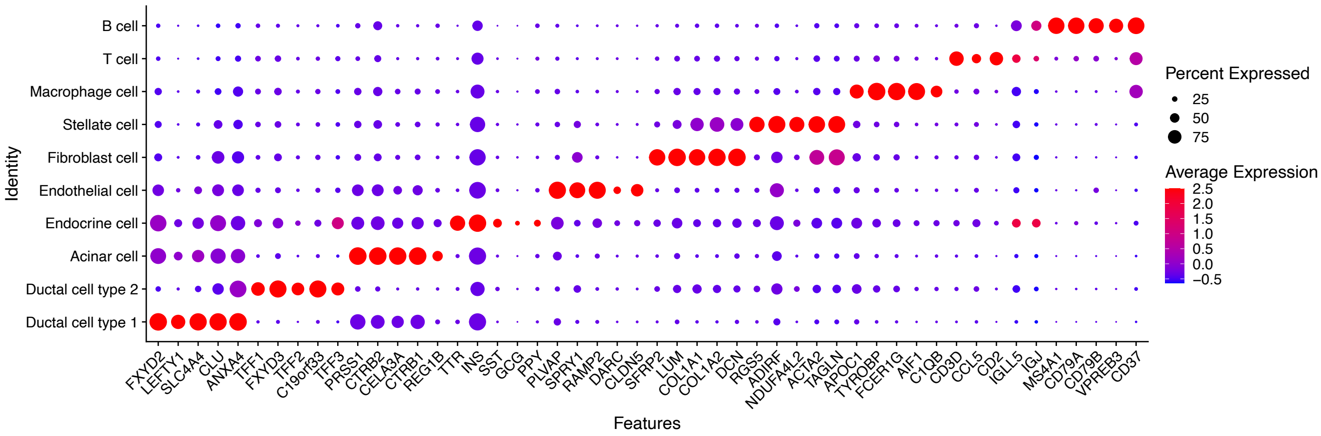


**Figure S12.** Markers used to characterize clusters in the PDAC single cell data set.

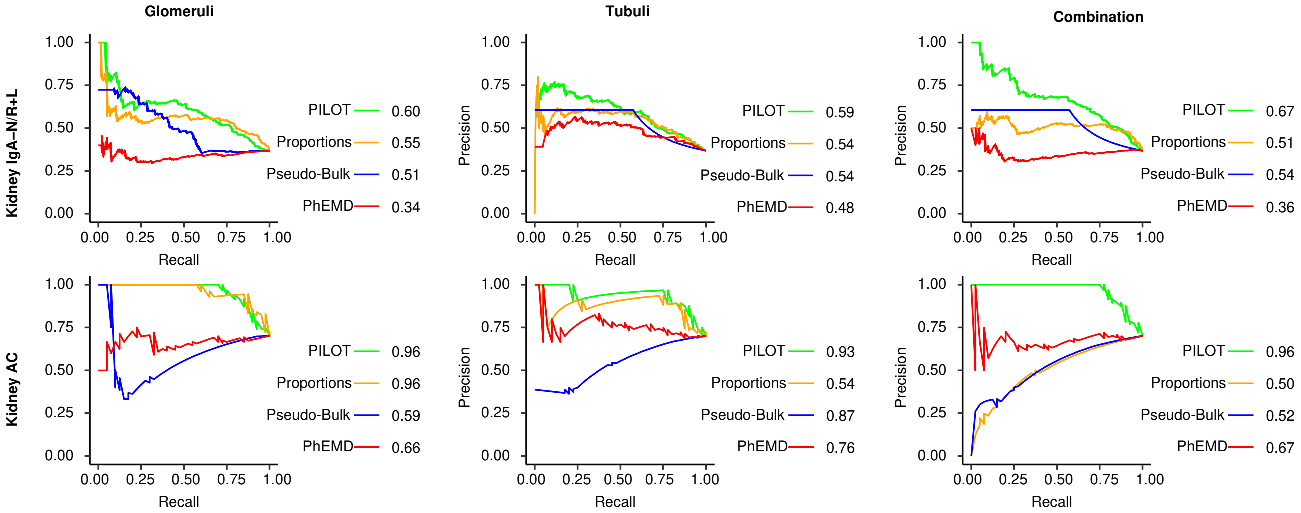


**Figure S9.** AUCPR plots for Glomeruli, Tubule and both (combined) for Kidney AC, Kidney IgAN pathomics data.

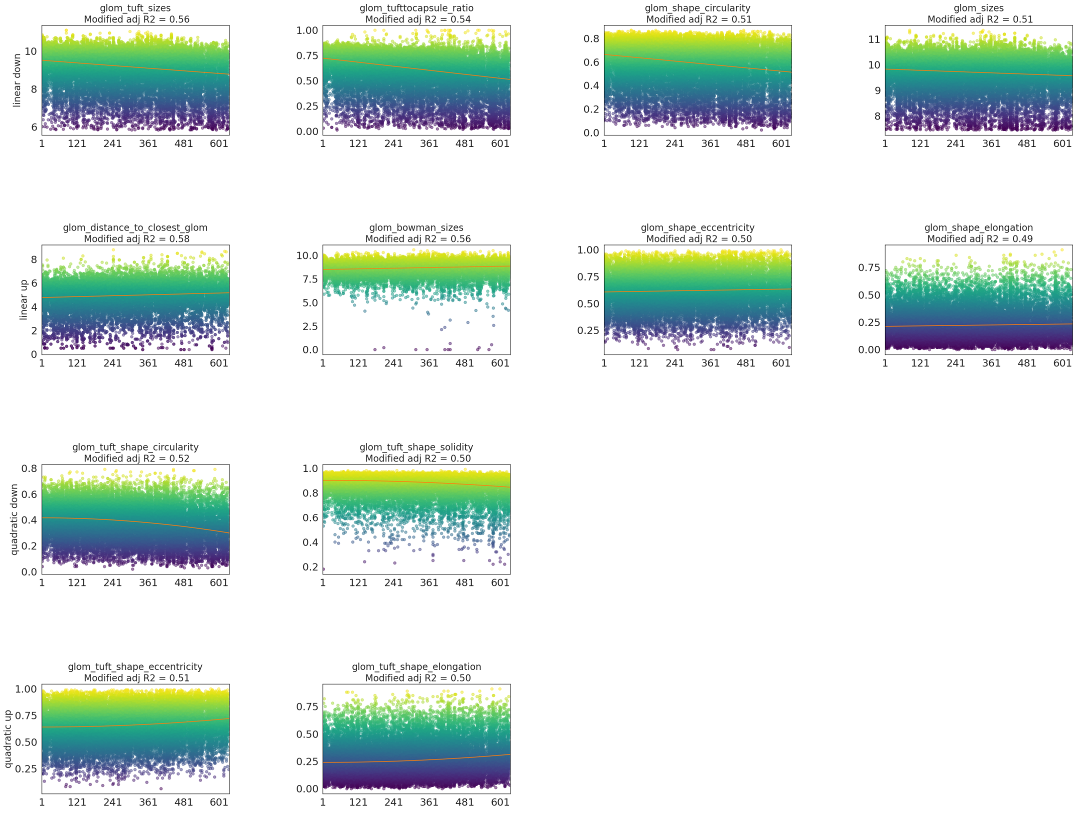


**Figure S10.** Top morphological features associated (ranked by adjusted R2) from Kidney IgA gromeruli. Every line corresponds to distinct models (linear, quadratic, linear-quadratic) and patterns (up vs. down). Only significant features ($p$-value > 0.01) are shown.

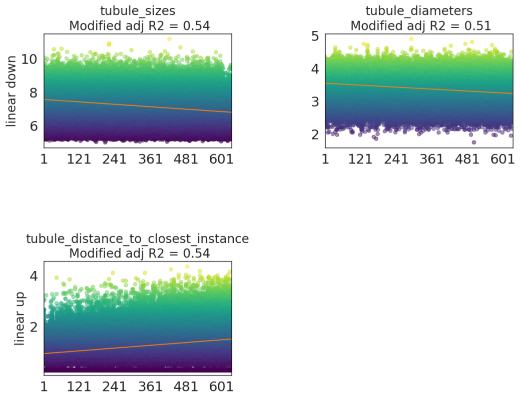


**Figure S11.** Top morphological features associated (ranked by adjusted R2) from Kidney IgA tubule. Every line corresponds to distinct models (linear, quadratic, linear-quadratic) and patterns (up vs. down). Only significant features (*p*-value > 0.01) are shown.
